# *Mx1-Cre*-mediated *Rac1* knockout confers multiple protective effects against anthracycline-induced acute normal tissue injury

**DOI:** 10.1101/2025.07.03.662930

**Authors:** Pelin Kücük, Sophie Gatzmanga, Jana Aengenvoort, Christian Henninger, Caroline Heinrich, Arthiga Vijayendran, Gerhard Fritz

**Affiliations:** Institute of Toxicology, Medical Faculty and University Hospital Düsseldorf, Heinrich Heine University Düsseldorf, Moorenstrasse 5, Düsseldorf 40225, Germany

**Author notes:** Corresponding authors: Prof. Dr. Gerhard Fritz, Institute of Toxicology, Medical Faculty and University Hospital Düsseldorf, Heinrich Heine University Düsseldorf, Moorenstrasse 5, 40225 Düsseldorf, Germany, Dr. Pelin Kücük, Institute of Toxicology, Medical Faculty and University Hospital Düsseldorf, Heinrich Heine University Düsseldorf, Moorenstrasse 5, 40225 Düsseldorf, Germany.

**Keywords:** RAC1, anticancer drugs, cardiotoxicity, hepatotoxicity, nephrotoxicity, DNA damage, apoptosis

## Abstract

Pharmacological data point to RAC1 as promising target for protection against anthracycline-induced cardiotoxicity, yet supporting genetic evidence is limited. Moreover, the relevance of RAC1 for cross-organ injury and cross-agent-induced normal tissue toxicity is unknown. Here, we employed a *Mx1-Cre*-based mouse model that enables an inducible *Rac1* knockout across different organs in order to comparatively analyze the influence of RAC1 on doxorubicin (DOX) and cisplatin (CisPt)-induced acute stress responses of the heart, kidney and liver. Following DOX treatment, the extend of DNA-double strand break (DSB) formation and the percentage of apoptotic cells were reduced in all three organs (i.e. heart, liver, kidney) of RAC1 deficient (Rac1^-/-^) animals as compared to the wildtype control (Rac1^+/+^). By contrast, in the absence of RAC1, CisPt-induced DNA damage was reduced only in the liver and the frequency of CisPt-stimulated apoptotic cell death remained unaffected by the Rac1 status in all organs. These findings demonstrate a strikingly organ- and agent-specific relevance of RAC1-regulated mechanisms for normal tissue damage evoked by genotoxic anticancer therapeutics. Accordingly, protein levels of phosphorylated DNA damage response (DDR)-related factors were also reduced in the kidney and liver of DOX treated Rac1^-/-^ animals, but not in the heart. Yet, DOX-triggered mRNA expression of surrogate markers related to inflammation, fibrosis and senescence was preferentially reduced in the heart and kidney of *Rac1* deficient mice. Taken together, RAC1 plays a so far unknown distinct role in the pathophysiology of anticancer drug-induced normal tissue damage by influencing DNA damage formation, activation of DDR, apoptosis induction as well as inflammation-, fibrosis- and senescence-associated responses in a pronounced agent- and organ-specific manner. Targeting of RAC1 appears particularly effective in the context of anthracycline-based therapeutic regimen for conferring broad organoprotection to both the heart and detoxifying organs.

**Translational Perspective:** This study provides the first genetic evidence identifying RAC1 as a critical mediator of anthracycline-induced cardiac, hepatic and renal injury. We demonstrate that genetic deletion of *Rac1* alleviates chemotherapy-induced DNA damage, apoptosis, and stress responses related to senescence, fibrosis, and inflammation in an agent- and organ-specific manner. These findings highlight RAC1 as a putatively clinically relevant target for the development of organoprotective pharmacological strategies aiming to improve the tolerability of anthracycline- and cisplatin-based therapeutic regimen and, accordingly, enhancing the quality of life and prognosis of tumor patients.

## 1. Introduction

Doxorubicin (DOX) is widely used in anticancer therapy (1). Its dose-limiting adverse effect is cardiotoxicity (2), which depends on the cumulative dose and manifests as congestive heart failure (CHF) due to cardiomyopathy (3). Cardiac damage can occur either at early time point after drug administration or emerge up to several years later (4). Although anthracyclines are well-known for causing cardiovascular damage, they also evoke substantial hepato- and nephrotoxicity (1, 5–7). The pathophysiological mechanisms suggested to be involved in anthracycline-induced normal tissue damage are highly complex and controversially discussed (8, 9). Among other, anthracyclines lead to the formation of DNA double-strand breaks (DSB) by topoisomerase II (TOP2) poisoning (10), especially of the β-isoform TOP2B (11–13). Since DSB are highly cytotoxic DNA lesions and potent activators of the DNA damage response (DDR), which defines the balance between survival and death on the molecular level (14–16), it is rational to assume that DSB contribute to the pathophysiology of anthracycline-induced heart damage (17). In addition to DSB, the generation of peroxynitrite by inducible nitric oxide synthases (iNOS), formation of reactive oxygen species (ROS), mitochondrial damage, DNA intercalation and histone eviction are also among the proposed pathophysiological mechanisms contributing to anthracycline toxicity (3, 17–19). It is plausible that the molecular mechanisms underlying organ toxicity depend on the anthracycline dose, treatment plan (single vs. repeated treatment) and time point of analyses (8). In line with this, it is assumed that acute and late toxicity evoked by anthracyclines involve distinct pathophysiological mechanisms (8, 20).

RAS-homologous (RHO) GTPases are well-known as key regulators of the actin cytoskeleton (21–23). However, they are also involved in the regulation of multiple other cellular functions, including gene expression, proliferation, inflammation and cell death (24–30). Noteworthy, RHO GTPases interfere with cardiovascular diseases (31, 32), including cardiac ischemia/reperfusion damage (33), myocardial hypertrophy (34) and atrial fibrillation (35), demonstrating their relevance for the regulation of heart functions. In addition, damage to the liver (36, 37) and kidney (38, 39) were also related to RHO signaling. HMG-CoA reductase inhibitors (statins), a class of cholesterol-lowering drugs that are well-known for their multiple beneficial effects on the cardiovascular system, counteract anthracycline-induced cardiotoxicity (40–42) and, moreover, mitigate hepatotoxicity (37) and kidney disease (43). The mechanism of statin action has been attributed to the inhibition of C-terminal prenylation of RHO GTPases, rather than to their cholesterol-lowering effect (44–46). Of note, among the different members of the family of RHO GTPases, signal mechanisms regulated by RAC1 (RAS-related C3 botulinum toxin substrate 1) GTPase are believed to be of major relevance for cardiac damage caused by anthracyclines (47–50). However, pinpointing RAC1 as the crucial target of statin-mediated cardioprotection is challenging, since statin-mediated inhibition of HMG-CoA reductase and resulting depletion of isoprene moieties can basically affect various types of RHO GTPases. For instance, radioprotective effects of statins have been mainly attributed to the inhibition of RHO/ROCK signaling (51, 52) rather than RAC1 signaling. On the other hand, frequently used small-molecule inhibitors of RAC1, such as NSC23766 and EHT1864, confer similar protective effects against DOX-induced cardiotoxicity *in vitro* and *in vivo* as statins (49, 53, 54), supporting the hypothesis that RAC1-regulated signaling is of particular relevance for cardiac injury following doxorubicin treatment. Notably yet, these inhibitors mitigate RAC1 activity by blocking the RAC1-GEF TIAM1 (i.e., NSC23766) or disrupting RAC1 effector coupling (i.e., EHT1864), respectively (55, 56). Thus, it cannot be ruled out that these compounds also affect the activity of other GTPases, such as CDC42 or RAC2/RAC3. Accordingly, the influence of different types of RHO GTPases on anthracycline-triggered cardiac injury remains obscure.

Although the aforementioned NSC23766- and EHT1864-based studies point to RAC1-regulated mechanisms as putative key players in anthracycline-induced cardiotoxicity, supportive genetic evidence for this hypothesis is largely missing but highly preferable. In this context, we previously showed that cardiomyocyte-specific *Rac1* knockout conferred only partial protection against anthracycline-induced cardiac damage (57), indicating that either other RHO GTPases than RAC1 and/or additional cell types contribute to anthracycline-induced cardiac damage. In line with this hypothesis, recently reported *in vitro* and *in vivo* data showed that non-cardiomyocyte cell types of the heart are comparably susceptible to anthracycline-induced injury as cardiomyocytes, involving both RAC1- and CDC42-regulated mechanisms (58). Importantly, statins have also been reported to protect against anthracycline-induced hepato-renal toxicity (59), arguing for a broader usefulness of this class of clinically well-established drugs to limit adverse effects of anthracyclines on multiple normal tissues. In addition, available data also point to a possible nephroprotective activity of statins against CisPt-induced nephrotoxicity (60, 61). This finding indicates that organoprotective effects of statins are not limited to adverse effects of anthracyclines only but may comprise different types of anticancer therapeutics. However, we also observed that an Alb-Cre mediated hepatic *Rac1* knockout protects the liver from DOX-induced acute and subacute stress responses but fails to confer protection against ionizing radiation-induced DNA damage (62). Thus, it cannot be ruled out that RAC1 preferentially contributes to anthracycline-induced stress responses while other RHO GTPases may be more relevant for the outcome of DNA damage caused by other types of genotoxic agents. For instance, transcript levels of both *Cdc42* and *Rho* are upregulated in proximal tubule kidney cells following CisPt treatment (63). Hence, hypothesizing that RAC1 is a biologically relevant target of statins regarding their presumed multiple beneficial effects on anticancer drug-induced normal tissue damage, it is tempting to speculate that RAC1 signaling influences multiple adverse outcome pathways of anticancer drugs in various organs in an agent- and tissue type-specific manner.

To substantiate this hypothesis by use of a genetic approach, we employed a Poly(I:C)-inducible *Rac1*^flox/flox^/*^Mx1-Cre^*mouse model in the present study. Thereby, a broad *Rac1* knockout in various responsive cell types and tissues is achieved (64, 65). Using this genetic model, we investigated the impact of RAC1 signaling on adverse outcome pathways triggered by different anticancer drugs (i.e., doxorubicin, cisplatin) in different organs (i.e., heart, liver, kidney). To this end, the steady state levels of DSB, the frequency of apoptosis and alterations in gene expression that are indicative of ongoing pathophysiological processes were comparatively investigated in RAC1 proficient (Rac1^+/+^) versus RAC1 deficient (Rac1^-/-^) mice.

## 2. Materials and Methods

### 2.1. Materials

Drugs, reagents and assay kits were purchased from following providers: Polyinosinic:polycytidylic acid (Poly(I:C)) (Sigma Aldrich, Darmstadt, Germany), DOX (Cellpharm, Bad Vilbel, Germany), CisPt (Teva, Petah Tikwa, Israel), medical dipsticks detecting GOT and GPT aminotransferases (Reflotron®, Roche Diagnostics Ltd., Rotkreuz, Switzerland), cTNI ELISA Kit (Life Diagnostics, West Chester, USA), KIM-1 ELISA Kit (RayBiotech, Peachtree Corners, USA), DNeasy Kit (Qiagen, Hilden, Germany), RNeasy Mini Kit (Qiagen, Hilden, Germany), High-Capacity cDNA Reverse Transcription Kit (Qiagen, Hilden, Germany), SensiMix SYBR Kit (Bioline Reagents Ltd., London, UK), TUNEL assay (Roche Diagnostics, Mannheim, Germany), DAPI containing antifade mounting medium (Thermo Fisher Scientific, Waltham, MA, USA), Alexa Fluor^TM^ 555-conjugated WGA (wheat germ agglutinin) (Sigma Aldrich, Darmstadt, Germany). Antibodies were purchased from following providers and used at indicated dilutions: anti-phospho-H2AX (Ser139) (1:500) (#9718S, Cell Signaling, Beverly, MA, USA), anti-alpha smooth muscle actin (#ab5694) (abcam, Cambridge, UK) (1:5000); anti-phospho-p53 (S15) (#9284); anti-GAPDH (#2118) (Cell Signaling, Beverly, MA, USA); anti-phospho-H2AX (Ser139) (#05-636); anti-Rac1 (#05-389), anti-PCNA (#MAB424R) (Merck Millipore, Darmstadt, Germany); anti-phospho-KAP1 (S824) (#A300-767A) (Bethyl Laboratories, Montgomery, TX, USA) (1:1000); HRP-conjugated secondary antibodies (#610-1302 or #611-1302, Rockland Immunochemicals Inc., Limerick, Pam USA) (1:2000); Alexa Fluor^TM^ 488-conjugated secondary antibody (#A11008, Invitrogen, Darmstadt, Germany) (1:600).

### 2.2. Animal experiments and treatment groups

Animal experiments were performed in accordance with the European Guidelines for the Care and Use of Laboratory Animals (2010/63/EU) and were approved by the North Rhine-Westphalia State Agency for Nature, Environment and Consumer Protection (approval reference number: 81-02.04.2020.A140). Mice were maintained in the central animal facility of the Heinrich Heine University Düsseldorf (Germany). 8-10 weeks old male and female mice (C57BL/6 background) were used at a group size of 4-8 animals per experimental group. *Rac1*^flox/flox/*Mx1-Cre*^ mice were treated with Poly(I:C) to induce *Rac1* knockout (KO) as described previously (65). *Rac1* wild-type (WT) (*Rac1*^flox/flox^) mice were also treated with Poly(I:C) as control. 3 weeks after the Poly(I:C) treatment, mice were injected with a single dose of DOX (24 mg/kg BW, i.p.) or saline. DOX-induced acute responses were analyzed after 48 h **(Fig. 1A)**. A single injection of CisPt (20 mg/kg BW, i.p.) was administered in a similar experimental setting for control of agent specificity **(Supplementary Fig. S1)**.

**Figure 1.**
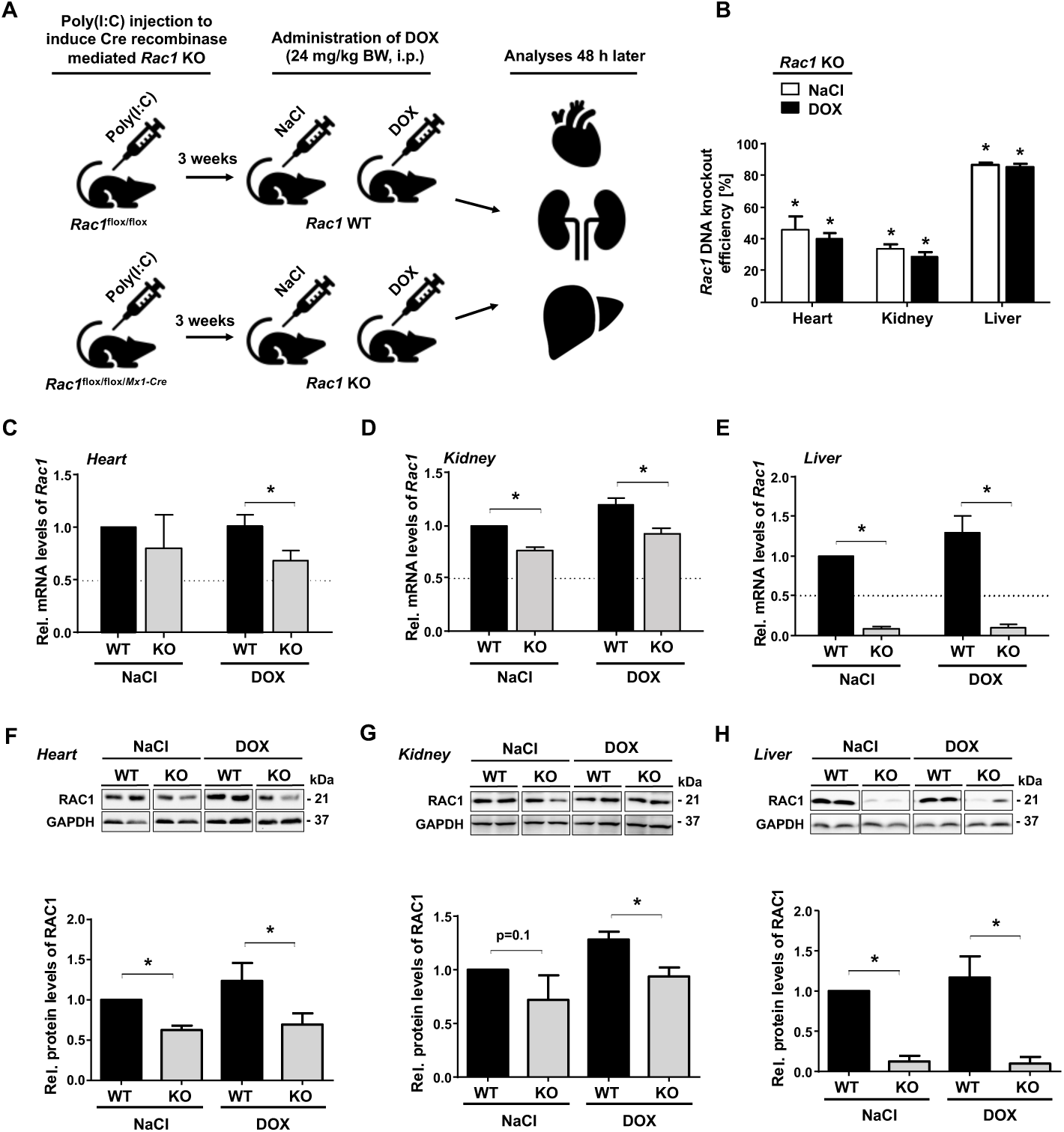
Generation of *Cre* recombinase-mediated *Rac1* knockout mice and DOX treatment regimen. **A)** Poly(I:C) was administered to *Rac1*^flox/flox/*Mx1-Cre*^ and *Rac1*^flox/flox^ mice to induce *Cre* recombinase-mediated *Rac1* knockout (KO) and to generate matched *Rac1* wild-type controls (WT), respectively. A single injection of DOX (24 mg/kg BW, i.p.) was administered 3 weeks later and analyses were performed after post-incubation period of 48 h. Control groups were treated with saline. **B)** *Rac1* DNA knockout efficiency was comparatively analyzed by genomic PCR in both the heart, kidney and liver. Data represent the mean + SEM from n=3 individual mice analyzed per experimental group. *p ≤ 0.05, one-tailed Student’s *t*-test. *vs corresponding wild-type controls. **C-E)** mRNA levels of *Rac1* were monitored by RT-qPCR in the heart **(C)**, kidney **(D)** and liver **(E)** tissues. mRNA expression levels were normalized to *Gapdh* and *Actb.* Relative *Rac1* mRNA expression in the saline-treated *Rac1* WT control group was set to 1.0 and relative *Rac1* mRNA expression levels are shown as the mean + SEM from n=3 mice per experimental group. *p ≤ 0.05, one-tailed Student’s *t*-test. **F-H)** RAC1 protein expression was investigated by Western blot analysis in the heart **(F)**, kidney **(G)** and liver **(H)**. GAPDH expression was used as protein loading control. Upper panels: representative Western blots; Lower panels: For quantitative evaluation, RAC1 protein expression in saline-treated RAC1 WT control group was set to 1.0 and relative RAC1 protein levels are shown as mean + SEM from n=3-6 mice per experimental group. *p ≤ 0.05, one-tailed Student’s *t*-test.

### 2.3. Analysis of blood parameters

Peripheral whole blood was collected into EDTA-coated tubes was analyzed for blood count using Scil Vet ABC Hematology Analyzer (Scil Animal Care Company, Illinois, USA). Blood samples were left at room temperature to allow coagulation and centrifuged at 10.000 ×g for 10 min to collect serum. Serum parameters were determined using a Reflotron® clinical chemistry analyzer and Reflotron® medical dipsticks detecting glutamate oxaloacetate transaminase (GOT) and glutamate pyruvate transaminase (GPT), indicative of liver injury. ELISA-based assays were used to detect serum cardiac troponin I (cTNI) and kidney injury molecule-1 (KIM-1), indicative of cardiac and kidney injury, respectively.

### 2.4. Isolation of DNA and RNA

Total DNA was purified from 5-10 mg of tissue using the TissueLyser II (Qiagen, Hilden, Germany) and DNeasy Kit from three mice per experimental group to assess *Rac1* gene knockout efficiency. Total RNA was purified from 20-30 mg of tissue using the TissueLyser II and the RNeasy Mini Kit for gene expression analyses. High-Capacity cDNA Reverse Transcription Kit (Thermo Fisher Scientific, Waltham, MA, USA) was used to perform reverse transcriptase reaction using 1000 ng of RNA per sample.

### 2.5. Real time RT-qPCR analysis

Real-time RT-qPCR analyses were performed using the SensiMix SYBR Kit in a thermal cycler (CFX96™ Real Time PCR Detection System (BioRad, Hercules, USA)). After initial denaturation at 95 °C for 10 min, 45 amplification cycles were performed (95 °C, 15 s; 55 °C, 17 s; 72 °C, 17 s), using corresponding gene-specific primers for DNA **(Supplementary Table 1)** and RNA **(Supplementary Table 2)**. Melting curves were analyzed to ensure product specificity. PCR products with threshold cycle (Ct) values ≥ 35 were excluded from analysis. mRNA expression levels were normalized to *Gapdh* and *Actb*. Gene expression of saline-treated WT animals was set to 1.0. RT-qPCR analyses were performed using pooled RNA samples isolated from three mice per experimental group, unless otherwise indicated. Changes in gene expression of ≤0.5-fold and ≥2-fold were considered as biologically relevant. Statistical analyses were not conducted when pooled RNA samples were used.

### 2.6. Immunohistochemistry and Immunofluorescence microscopy

Formalin-fixed, paraffin-embedded heart, liver and kidney tissues were cut into 3 μm thick sections using a Hyrax M25 microtome (Carl Zeiss, Jena, Germany). Paraffin removal, rehydration and antigen retrieval were performed according to standard procedure. Sections were incubated with Protein Block (Dako, Hamburg, Germany) (2 h; RT), followed by incubation with the corresponding primary antibody (overnight; 4 °C) and Alexa Fluor^TM^ 488-conjugated secondary antibody (2 h; RT). Alexa Fluor^TM^ 555-conjugated wheat germ agglutinin (WGA) (10 μg/ml) was used to stain cell membranes in the heart sections to allow discrimination between cardiomyocytes and non-cardiomyocytes and in the kidney sections to distinguish cortex, glomeruli and medulla. Sections were mounted using DAPI (Thermo Fisher Scientific, Waltham, MA, USA) containing antifade mounting medium and microscopic analysis was performed using an Olympus BX 43 microscope (Olympus, Hamburg, Germany).

### 2.7. Detection of DNA double-strand breaks (DSB) by nuclear **γ**H2AX foci analysis

DSB were monitored by immunofluorescence-based analysis of nuclear foci formed by Ser139-phosphorylated histone H2AX (γH2AX) as a surrogate marker of DSB (66), using anti-phospho-H2AX (Ser139) antibody. The percentage of γH2AX-positive cells was calculated following microscopic analysis using ImageJ software (67).

### 2.8. Detection of apoptotic cells by the TUNEL assay

Apoptotic cells were detected by the TUNEL assay according to manufacturer’s instructions. Briefly, free 3’-OH groups in fragmented DNA in apoptotic cells were labelled with fluorescein-conjugated dUTP nucleotides. The percentage of TUNEL-positive cells was calculated by microscopic analyses using the ImageJ software (67).

### 2.9. Detection of proliferating cells PCNA staining

Proliferating cells were detected by immunohistochemical analysis of proliferating cell nuclear antigen (PCNA). The percentage of PCNA-positive cells was calculated by microscopic analysis using ImageJ software.

### 2.10. Western blot analysis

15-20 mg of frozen tissue sections were homogenized in Roti^®^-Load 1 (Roth, Karlsruhe, Germany) using TissueLyser II (Qiagen, Hilden, Germany). Tissue lysate was sonicated and centrifuged (10 min at 10.000 ×g; 4 °C). The collected supernatant was heated for denaturation (95 °C, 5 min). Protein samples were separated by SDS-PAGE and transferred onto nitrocellulose membranes (GE Healthcare, Freiburg, Germany). After blocking (5% BSA, 0.1% Tween 20 in TBS), membranes were incubated with primary antibodies and subsequently with HRP-conjugated secondary antibodies. Protein expression of GAPDH was used as loading control. Chemiluminescence imaging was performed using the ChemiDoc™ Imaging System (Bio-Rad Laboratories, Hercules, CA, USA). For quantitation, densitometrical analysis was performed using ImageJ software.

### 2.11. Statistical analyses

For statistical analyses, Student’s *t*-test (unpaired, one- or two-tailed) was used (GraphPad Prism v6 (San Diego, CA, USA)). *p*-values ≤0.05 were considered as statistically significant differences and are marked with an asterisk (**\***).

## 3. Results

### 3.1. Generation of *Mx1-Cre* mediated *Rac1* knockout mice

We used a Poly(I:C) inducible *Mx1-Cre-*based mouse model (*Rac1*^flox/flox/*Mx1-Cre*^), which contains *LoxP* sites flanking *Rac1*, to generate *Rac1* knockout mice (65). Following a 3 weeks recovery period after Poly(I:C) induced *Cre*-recombinase expression, a single dose of DOX (24 mg/kg BW, i.p.) was administered. After a post-incubation period of 48 hours, the acute DOX-stimulated stress responses of the heart, kidney and liver were analyzed (**Fig. 1A)**. On the level of the genomic DNA, we detected ∼40%, ∼35% and ∼80% *Rac1* knockout efficiency in the heart, kidney and liver, respectively **(Fig. 1B)**. Organ-specific differences in the *Rac1* knockout efficiency were further validated by analyzing *rac1* mRNA expression and RAC1 protein levels by RT-qPCR **(Fig. 1C-E)** and Western blot analyses **(Fig. 1F-H)**, respectively. In line with the genomic data the results of these analyses revealed the most pronounced knockout of *Rac1* in the liver.

### 3.2. Effect of *Rac1* knockout on body weight, blood parameters, and organ toxicity markers under basal situation and following DOX treatment

A significant loss of body weight was detected in both *Rac1* wild-type and *Rac1* knockout mice after DOX treatment as well as of *Rac1* knockout mice under basal situation **(Fig. 2A)**. The relative weight of the heart and kidney also decreased after DOX exposure **(Fig. 2B-C)**, while the relative liver weight remained unchanged **(Fig. 2D)**. The *Rac1* status neither influenced body nor organ weight **(Fig. 2A-D)**. DOX treatment significantly reduced leukocyte counts in both *Rac1*^+/+^ and Rac1^-/-^ animals **(Fig. 2E)**. A tendential increase in monocyte counts, as well as a significant increase in neutrophil counts were detected after DOX treatment in wild-type mice, which was not detectable in the absence of *Rac1* **(Fig. 2E)**. Thus, *Rac1* deficiency prevents DOX-stimulated increase in neutrophils indicating that RAC1 may contribute to the inflammatory response triggered by DOX. We also evaluated the platelet to lymphocyte ratio as a prognostic marker for cardiac damage (68, 69). DOX led to a marked increase in the platelet to lymphocyte ratio in both *Rac1* wild-type and knockout animals **(Fig. 2F)**. Analyzing cardiac troponin I (cTNI) kidney injury molecule (KIM-1) or transaminases (GOT and GPT), which are markers for cardiotoxicity, nephrotoxicity and hepatotoxicity, respectively, revealed no significant changes at the time point of analysis **(Supplementary Fig. S2)**. However, measuring RNA levels of *Nppa* and *Nppb* in the heart tissue as markers of cardiac damage, we detected a significant increase in *Nppa* mRNA levels evoked by DOX, which was tendentially reduced in *Rac1* knockout animals **(Fig. 2G)**. Elevated *Nppb* mRNA levels were observed under basal situation in saline-treated *Rac1* knockout animals but did not further increase following DOX treatment **(Fig. 2G)**. Regarding kidney toxicity, DOX treatment did not affect *Kim-1* mRNA expression levels **(Fig. 2H)**. In contrast, *Gstm1* mRNA levels were markedly increased in the liver of DOX-treated wild-type animals. This effect, which is indicative of the activation of hepatic stress responses, was not observed in *Rac1* knockout animals **(Fig. 2I)**, indicating that RAC1 deficiency has a hepatoprotective effect. In summary, DOX-induced stress responses are organ specific and differently affected by the *Rac1* status.

**Figure 2.**
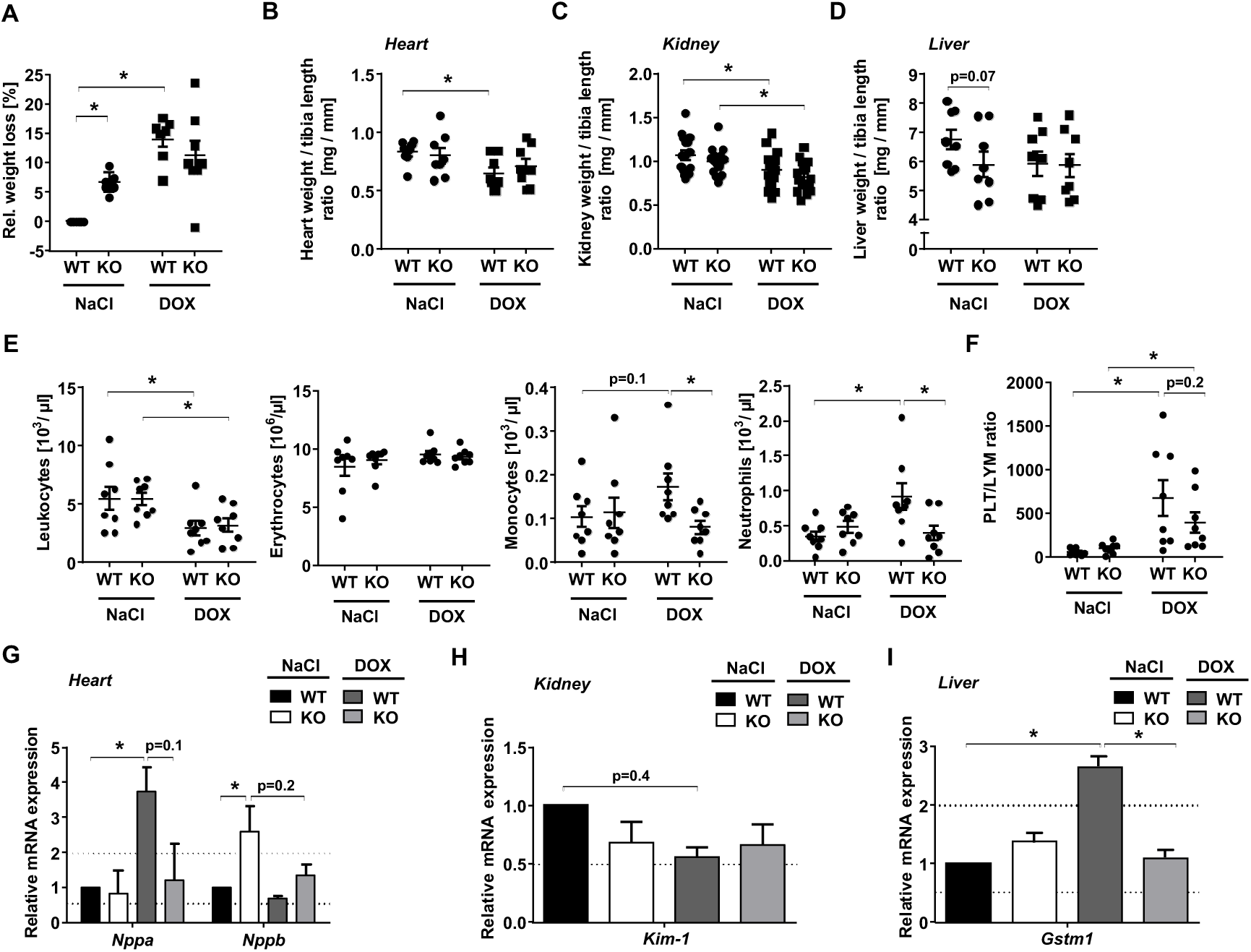
Influence of DOX treatment on body weight, blood parameters as well as cardiotoxicity, nephrotoxicity and hepatotoxicity markers in *Rac1* WT and *Rac1* KO mice. *Rac1* WT and *Rac1* KO mice were treated with saline or DOX as described above (Fig. 1A). **A)** Weight loss is shown in percentage of total body weight at the end of the experiment. Data obtained for each mouse are shown as well as the mean ± SEM. *p≤0.05, two-tailed Student’s *t*-test. **B-D)** Organ weight was calculated relative to tibia length in the heart **(B)**, right and left kidneys **(C)** and liver **(D)**. Data obtained for each mouse as well the mean ± SEM are shown. *p≤0.05, two-tailed Student’s *t*-test. **E)** Blood samples were analyzed for leukocyte, erythrocytes, monocyte and neutrophil counts and counts obtained for each mouse as well the mean ± SEM are presented. *p≤0.05, two-tailed Student’s *t*-test. **F)** Platelet-lymphocyte ratio was calculated. Data are shown as the mean ± SEM from n=7-8 mice per experimental group. *p≤0.05, two-tailed Student’s *t*-test. **G-I)** mRNA expression levels of organ specific toxicity-related markers *Nppa* and *Nppb* **(G)**, *Kim-1* **(H)** and *Gstm1* **(I)** were analyzed in the heart, kidney and liver tissue, respectively. mRNA expression levels were normalized to *Gapdh* and *Actb.* Relative mRNA expression in saline-treated *Rac1* WT control group was set to 1.0 and relative mRNA expression levels are shown as the mean + SEM from n=3-4 mice per experimental group. *p ≤ 0.05, two-tailed Student’s *t*-test.

### 3.3. *Rac1* knockout protects heart, kidney and liver tissues from DOX-induced DNA damage formation

To investigate the level of DOX-induced DNA damage (i.e., DSB), the percentage of γH2AX foci positive cells was determined in tissue sections of the heart, kidney and liver **(Fig. 3A-C)**. Alexa Fluor™-conjugated WGA staining was used to distinguish between different cell types in the heart (i.e., cardiomyocytes and non-cardiomyocytes) and between different anatomical regions in the kidney (i.e., cortex, medulla and glomeruli). Under basal situation we observed that the percentage of γH2AX-positive cells was relatively high (∼20%) in cardiomyocytes of saline-treated *Rac1* wild-type and knockout animals as compared to non-cardiomyocytes (∼3%) **(Fig. 3A)**. DOX treatment led to a significant increase in the percentage of γH2AX-positive cells, both in cardiomyocytes (∼3-fold) and non-cardiomyocytes (∼5-fold) of WT mice **(Fig. 3A)**. Importantly, DOX-induced DNA damage was significantly attenuated in both cardiac cell populations in the *Rac1* knockout mice as compared to the wildtype animals **(Fig. 3A)**. This finding demonstrates that Rac1 deficiency confers substantial genoprotection against DOX-induced DNA damage across different cardiac cell types. In the kidney, basal levels of γH2AX-positive cells were similar in cortex and medulla (∼5%). DOX treatment significantly increased the percentage of γH2AX-positive cells (∼8-fold) in both cortex and medulla **(Fig. 3B)**. Like in the heart, Rac1 knockout significantly reduced DOX-induced DNA damage in both cortical and medullary kidney cells **(Fig. 3B)**. On the contrary, DOX-evoked increase in the percentage of γH2AX-positive glomerular cells remained unaffected by *Rac1* knockout **(Fig. 3B)**, highlighting cell type-specific differences. The most pronounced increase in the percentage of γH2AX-positive cells after DOX treatment was observed in the liver of WT animals (∼30-fold as compared to untreated control), which was almost entirely prevented in the *Rac1* knockout animals **(Fig. 3C)**. This finding demonstrates a substantial protection against DOX-induced hepatotoxicity resulting from *Rac1* knockout in the liver.

**Figure 3.**
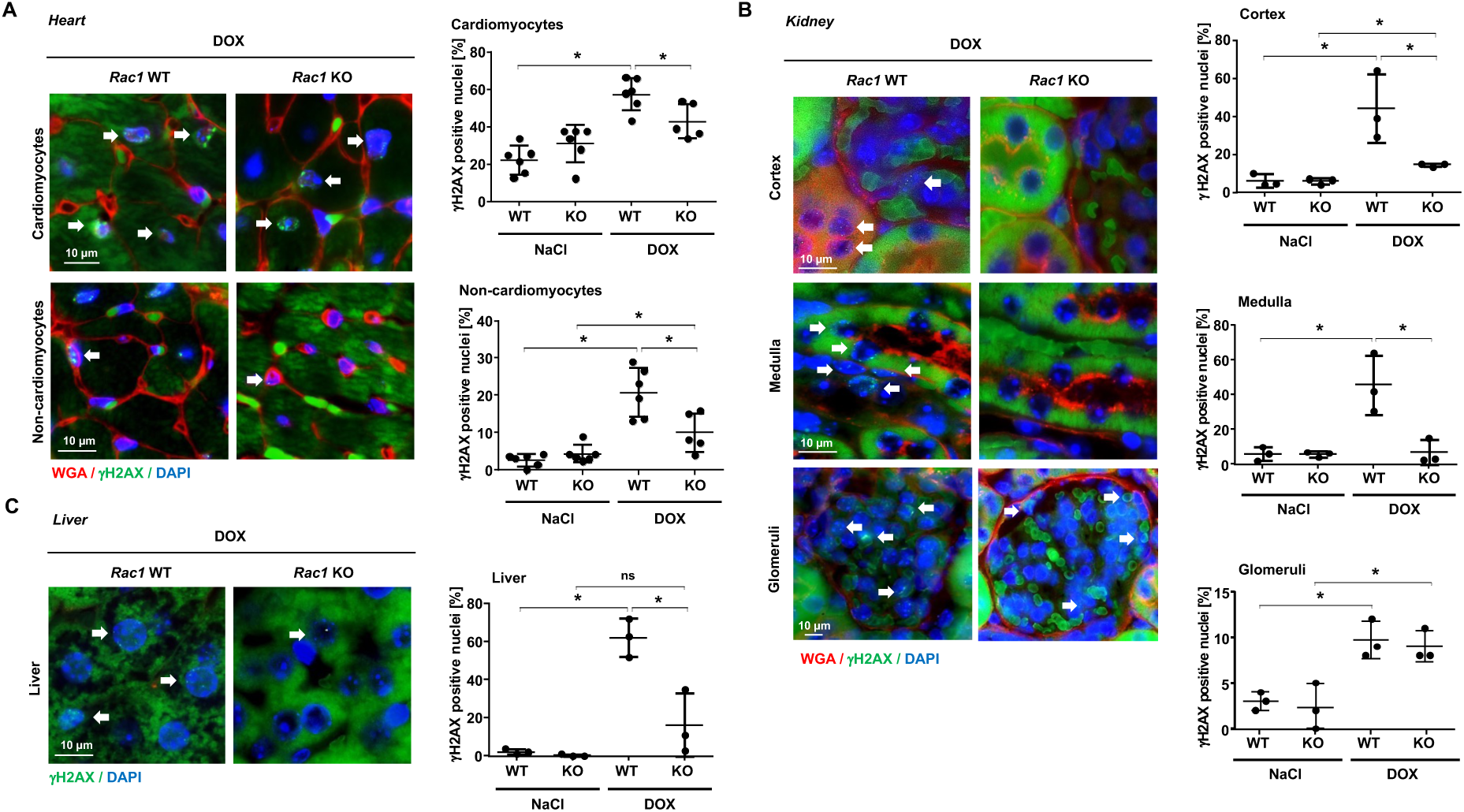
Influence of *Rac1* knockout on DOX-induced DSB levels in the heart, kidney and liver. *Rac1* WT and *Rac1* KO mice were treated with saline or DOX as shown before (Fig. 1A). **A-C)** DNA DSB were visualized by immunohistochemical staining of γH2AX foci in tissue sections of the heart **(A)**, kidney **(B)** and liver **(C)**. WGA staining was used to visualize cell borders **(A)** in order to distinguish between cardiomyocytes and non-cardiomyocytes in the heart and **(B)** cortex, medulla and glomeruli in the kidney. The percentage of γH2AX positive nuclei was quantified by microscopic analysis. Quantitative data shown in the histograms are the mean ± SD from n=3-6 animals per experimental group with two technical replicates and >150 nuclei being analyzed per experimental condition. *p ≤ 0.05; ns, not significant; two-tailed Student’s *t*-test. The left panels show representative pictures. WGA (red), γH2AX (green), DAPI (blue); 100x objective. White arrows indicate γH2AX positive nuclei. Representative pictures from control groups are shown in **Supplementary Fig. S4**.

To determine whether *Rac1* knockout mediates drug-specific organoprotection, we analyzed organ-specific stress responses following treatment of the mice with the platinating anticancer drug cisplatin (CisPt). CisPt also increased the proportion of γH2AX-positive cells in the heart, kidney and liver **(Supplementary Fig. S3A-C)**. However, protection against CisPt-induced DNA damage by *Rac1* knockout was observed only in the liver **(Supplementary Fig. S3C)**, demonstrating cross-agent hepatoprotective effects resulting from RAC1 deficiency. Importantly, *Rac1* knockout neither protected the heart nor the kidney **(Supplementary Fig. S3A-B)** from CisPt-induced DNA damage, further emphasizing organ-specific effects. Overall, our findings demonstrate that the RAC1 status has a distinct influence on the steady state level of anticancer drug-induced DNA damage in an organ-, tissue- and agent-specific manner. *Rac1* knockout mediates a pronounced genoprotection against DNA damage induced by DOX in multiple organs and tissues, while selectively protecting the liver against CisPt-induced DNA damage. Hence, we speculate that, specifically in the liver, a major proportion of DNA damage originates from a common pathway that is induced by both DOX and CisPt, which can be attenuated by *Rac1* deletion. In this context, it was previously reported that RAC1-signaling contributes to DOX-induced cardiotoxicity through both ROS-dependent and ROS-independent mechanisms (48).

### 3.4. *Rac1* knockout mitigated DOX-induced DNA damage response (DDR), particularly in the kidney and liver

Having in mind that DSB are potent triggers of the DDR, we next analyzed the impact of RAC1 on the ATM/ATR-mediated phosphorylation of representative DDR-associated proteins. DOX treatment resulted in a significant increase in protein levels of the chromatin regulatory factor pKAP1 in the heart, kidney and liver **(Fig. 4A-C)**. This effect was mitigated by *Rac1* knockout in the kidney and liver **(Fig. 4B-C)** but not in the heart **(Fig 4A)**. Similarly, γH2AX protein levels were elevated after DOX treatment in all organs analyzed **(Fig. 4A-C)**. *Rac1* knockout significantly reduced the DOX-induced increase in γH2AX protein levels in the liver **(Fig 4C)** and tendentially in the kidney **(Fig. 4B)** but not in the heart **(Fig. 4A)**. Of note, an increase in the basal level of γH2AX was observed in the heart of *Rac1* knockout animals **(Fig. 4A)**, which was not evident in the other organs. A DOX-induced marked increase in pP53 levels was specifically detected in the kidney, which was prevented by *Rac1* knockout **(Fig. 4B)**. In the heart **(Fig. 4A)** and liver **(Fig. 4C)**, DOX treatment slightly enhanced pP53 levels, which remained unaffected by *Rac1* knockout **(Fig. 4A and 4C)**. Of note, basal pP53 protein levels in the heart **(Fig. 4A)** and kidney **(Fig. 4B)** were significantly higher in *Rac1* knockout animals than in the wild-type control group. The findings demonstrate that RAC1 exerts organ-specific effects under basal conditions and, moreover, also modulates anthracycline-induced DDR-related mechanisms, including P53-related functions (50) that are pivotal in the regulation of DDR (16, 70), in an organ-specific manner.

**Figure 4.**
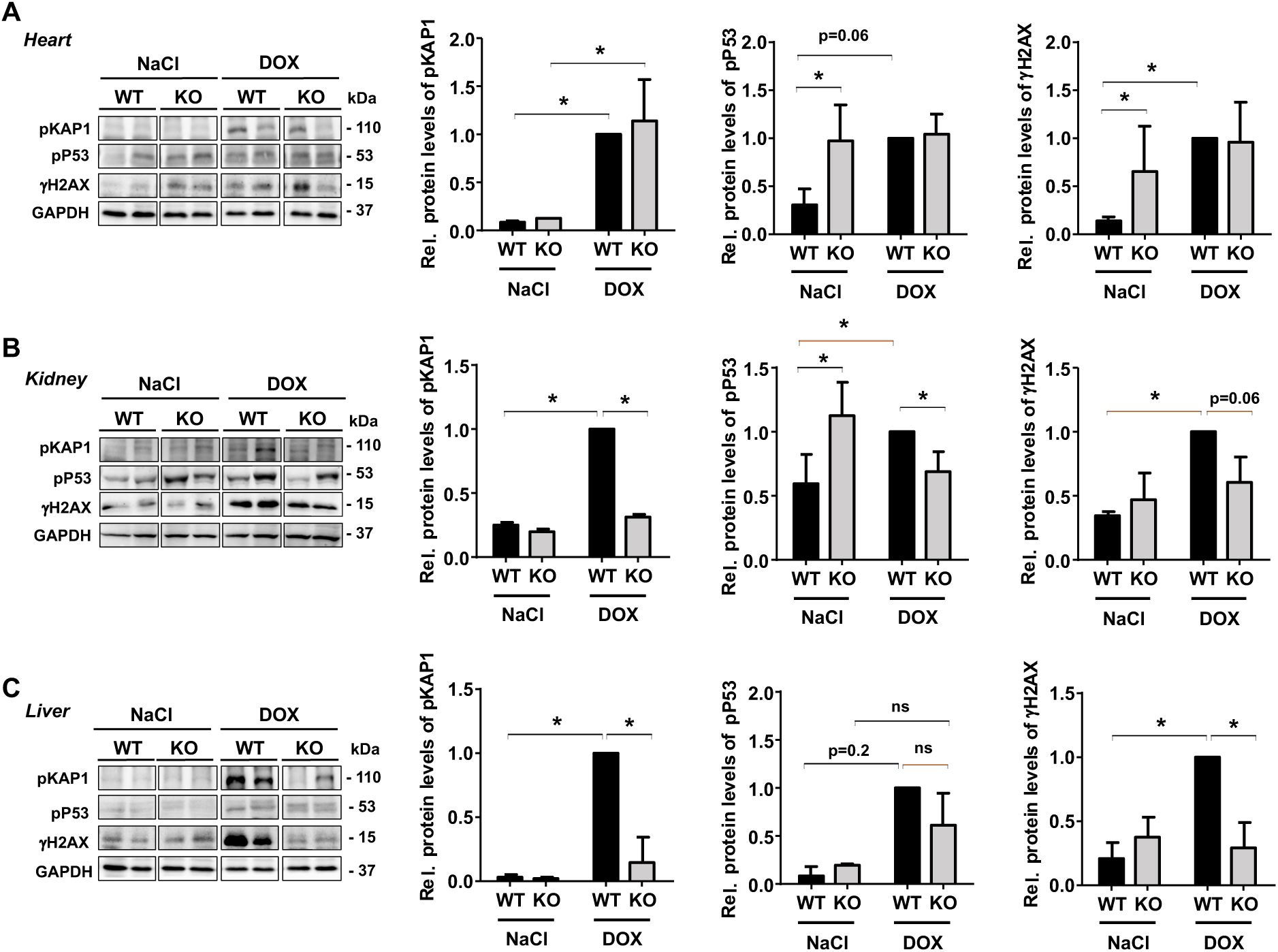
Influence of *Rac1* knockout on mechanisms of the DDR in the heart, kidney and liver. *Rac1* WT and *Rac1* KO mice were treated with saline or DOX as shown in Fig. 1A. **A-C)** Western blot analyses were performed to detect the protein expression levels of prototypical DDR-related markers γH2AX (S139), pP53 (S15) and pKAP1 (S824) in the heart **(A)**, kidney **(B)** and liver **(C)**. GAPDH was used as loading control. Protein expression in DOX-treated *Rac1* WT group was set to 1.0 and relative protein levels are shown as mean + SD from n=3-4 mice per experimental group. *p ≤ 0.05; ns, not significant; two-tailed Student’s *t*-test.

### 3.5. *Rac1* knockout protected the heart, kidney and liver tissues from DOX-induced apoptosis

Next, we investigated the influence of RAC1 on DOX-induced apoptosis by use of the TUNEL assay. DOX treatment triggered apoptosis only in non-cardiomyocytes, but not in cardiomyocytes of the heart *in vivo* **(Fig. 5A)**, which is in line with previously reported *in vitro* data (58). The percentage of DOX-induced TUNEL-positive non-cardiomyocytes was significantly reduced in the absence of *Rac1* **(Fig. 5A)**. Consistently, DOX-stimulated increase in the mRNA expression of the proapoptotic genes *Bax*, *Fasr* and *Fasl* in the heart was slightly diminished in *Rac1* knockout mice **(Fig. 5D)**. In the kidney, DOX treatment significantly enhanced the percentage of TUNEL-positive cells in both cortex and medulla, which was not observed in the *Rac1* knockout **(Fig. 5B)**. Among the apoptosis-related genes, DOX treatment enhanced only the mRNA expression of *Fasl* in the kidney, which was also diminished in the absence of *Rac1* **(Fig. 5E)**. Likewise, DOX led to a profound increase in the fraction of TUNEL-positive cells in the liver, which again was prevented by *Rac1* knockout **(Fig. 5C)**. DOX-stimulated increase in the hepatic mRNA levels of proapoptotic genes *Bax* and *Fasr* also decreased in the absence of *Rac1* **(Fig. 5F)**. The data demonstrate that the lack of RAC1 results in a distinct protection from DOX-induced apoptosis across different organs and cell types. Notably, CisPt also caused a substantial increase in the frequency of apoptotic cells in the heart, kidney and liver **(Supplementary Fig. S5A-C)**. However, *Rac1* knockout did not protect any of the organs against CisPt-induced apoptosis **(Supplementary Fig. S5A-C)**. Of note, the CisPt-induced apoptotic fraction was even further elevated in the liver if RAC1 is missing **(Supplementary Fig S5C)**. CisPt led to a substantial upregulation in the mRNA expression of apoptosis-related markers, including *Fasl* in the heart, *Bax* and *Fasr* in the kidney, and *Bax* in the liver. Among these, only the CisPt-induced increase in *Fasl* expression in the heart was attenuated in *Rac1* knockout animals, while the mRNA expression levels of the other markers remained unaffected by the *Rac1* status **(Supplementary Fig S5D-F).** These data highlight the agent-specific influence of RAC1 on anticancer drug-induced apoptotic mechanisms *in vivo* and, furthermore, demonstrate that *Rac1* knockout protects against DOX-induced apoptosis across different organs and cell types.

**Figure 5.**
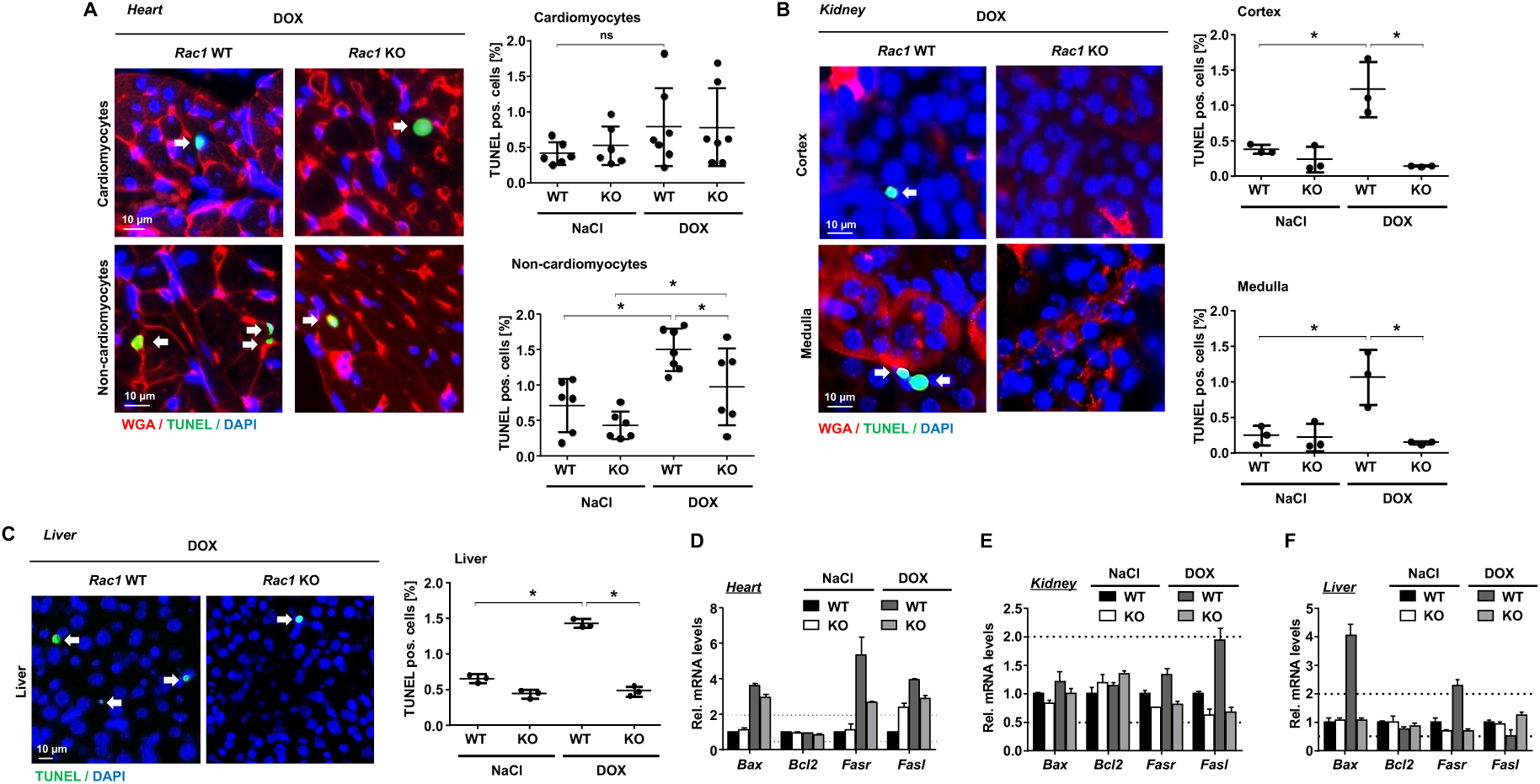
Influence of *Rac1* knockout on DOX-induced apoptosis in the heart, kidney and liver. *Rac1* WT and *Rac1* KO mice were treated with saline or DOX as shown in Fig. 1A. **A-C)** Apoptotic cells were detected by immunohistochemical TUNEL staining in the heart **(A)**, kidney **(B)** and liver **(C)** tissue sections. WGA was used to visualize cell borders in the heart **(A)** enabling the distinction between cardiomyocytes and non-cardiomyocytes and in the kidney **(B)** to distinguish between cells in the cortex and medulla. The percentage of TUNEL positive cells was quantified. Data shown are the mean ± SD from n=3-7 animals per experimental group, with two technical replicates and >300 cells being analyzed per condition. *p ≤ 0.05; ns, not significant; two-tailed Student’s *t*-test. The left panels show representative pictures. WGA (red), TUNEL (green), DAPI (blue); 100x objective. White arrows indicate TUNEL positive cells. Representative images from control groups are provided in **Supplementary Fig. S6**. **D-F)** mRNA expression levels of genes involved in the regulation of apoptosis were detected by RT-qPCR in the heart **(D)**, kidney **(E)** and liver **(F)** tissues. mRNA expression levels were normalized to those of *Gapdh* and *Actb.* Relative mRNA expression in saline-treated *Rac1* WT control group was set to 1.0 and relative mRNA levels are shown as mean + SEM from triplicate determination using pooled RNA samples isolated from n=3 animals per experimental group.

### 3.6. *Rac1* knockout modulated DOX-induced pathophysiological responses in the heart, kidney and liver

Next, we aimed to determine whether RAC1 influences tissue-specific responses to DOX-mediated injury. Using PCNA immunostaining, we assessed the proliferation index as a surrogate marker of tissue remodeling. Among the organs analyzed, only the heart showed a significant increase in the proportion of PCNA-positive cells following DOX treatment, which was further enhanced in the *Rac1* knockout animals **(Fig. 6A)**. In the kidney and liver, PCNA-positive fraction remained unchanged after DOX treatment **(Fig. 6B and 6C)**. Next, we investigated α-smooth muscle actin (αSMA) protein levels, as surrogate marker of fibrotic alterations, by Western blot analysis. Under basal conditions, RAC1 deficiency led to a significant increase in αSMA protein levels in the heart **(Fig. 6D)** and liver **(Fig. 6F)**, but not in the kidney **(Fig. 6E)**. Under our experimental conditions, DOX treatment had no significant influence on αSMA protein levels in the kidney and liver **(Fig. 6D, 6E and 6F)**. Measuring mRNA expression of markers indicative of ongoing senescence, fibrosis and inflammation, DOX treatment elevated the mRNA expression of *Cdkn1a*, *Ctgf*, *Il6* and *Mmp3* in the heart **(Fig. 6G)**. *Rac1* knockout prevented these cardiac stress responses resulting from DOX treatment **(Fig. 6G)**. In the kidney, DOX evoked a slight increase in *Cd19, Il2ra* and *Il6* mRNA levels. Among these, only the increase in *Cd19* mRNA levels were prevented by *Rac1* knockout **(Fig. 6H)**. In the liver, *Cdkn1a*, *Icam1* and *Il6* mRNA levels were enhanced in DOX treated animals. This increase remained largely unaffected by *Rac1* knockout, except for a reduction in *Icam1* mRNA expression **(Fig. 6I)**. Notably, basal *Cdkn1a* levels in the heart and liver of saline-treated *Rac1* knockout animals were relatively higher than in the wild-type control group **(Fig. 6G and 6I)**, while *Rac1* knockout reduced the mRNA levels of *Cdkn1a* under basal conditions in the kidney **(Fig. 6H)**. Collectively, the most pronounced pathophysiological tissue responses triggered by DOX were observed in the heart, which were largely diminished by *Rac1* knockout, further supporting the organ-specific role of RAC1 in modulating DOX-induced stress responses. Noteworthy, beyond its effect on DOX-stimulated stress responses, *Rac1* knockout differently influenced the basal mRNA expression of *Cdkn1a* and *Il6* in an organ-specific manner **(Fig 6G-I)**, indicating that RAC1 already interferes with mechanisms regulated by these factors under non-stressed conditions.

**Figure 6.**
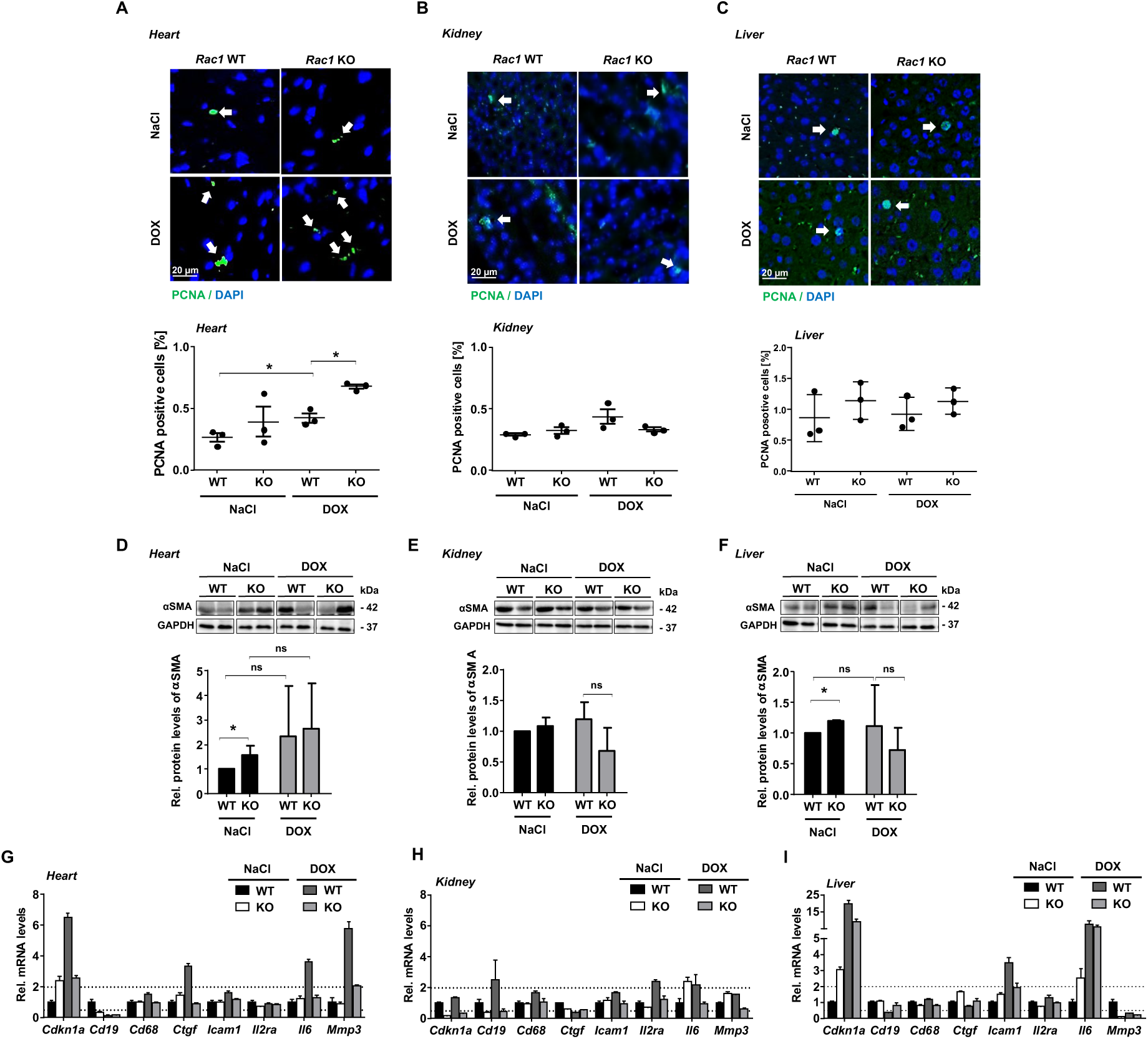
Influence of *Rac1* knockout on DOX-induced inflammation, fibrosis and senescence-related changes in the heart, kidney and liver. *Rac1* WT and *Rac1* KO mice were treated with saline or DOX as shown in Fig. 1A. **A-C)** Proliferative index was assessed by immunohistochemical PCNA staining in the heart **(A)**, kidney **(B)** and liver **(C)**. The percentage of PCNA-positive cells was quantified. Data shown are the mean ± SD from n=3 animals per experimental group with two technical replicates and >3000 cells being analyzed per condition. *p ≤ 0.05, two-tailed Student’s *t*-test. The upper panels show representative pictures. PCNA (green), DAPI (blue); 20x objective. White arrows indicate PCNA-positive cells. **D-F)** Western blot analysis was performed to analyzed protein levels of αSMA in the heart **(D)**, kidney **(E)** and liver **(F)**. GAPDH was used as loading control. αSMA protein expression in saline-treated *Rac1* WT control group was set to 1.0 and relative αSMA protein levels are shown as mean + SD from n=3-5 mice per experimental group. *p ≤ 0.05, two-tailed Student’s *t*-test. **G-I)** mRNA expression levels of selected genes involved in the regulation of senescence (*Cdkn1a*), inflammation (*Cd19*, *Cd68*, *Icam1*, *Il2ra*, *Il6*) or fibrosis (*Ctgf*, *Mmp3*) were investigated by RT-qPCR in the heart **(G)**, kidney **(H)** and liver **(I)** tissues. mRNA expression levels were normalized to *Gapdh* and *Actb.* Relative mRNA expression in saline-treated *Rac1* WT control group was set to 1.0 and relative mRNA levels are shown as mean + SEM from triplicate determinations using pooled RNA samples isolated from n=3 animals per experimental group.

In addition, we comparatively analyzed the basal and DOX-stimulated mRNA expression of a selected subset of genes involved in the regulation of RHO GTPase signaling, DNA repair, cell death, oxidative metabolism, mitochondrial homeostasis, topoisomerase function, senescence, inflammation and fibrosis across the different organs. Among the genes analyzed, the most pronounced changes were observed for *Cdh1* (E-cadherin) mRNA expression in the heart and *Acta2* (α-SMA) levels in the liver, indicative of tissue remodeling being influenced by DOX treatment and the *Rac1* status of the animals **(Supplementary Fig. S7).** Regarding CisPt-induced changes in the mRNA levels of senescence-, fibrosis- and inflammation-associated markers, CisPt treatment led to increased mRNA levels of *Icam1* and *Il6* in the heart. Notably, this upregulation was attenuated by *Rac1* knockout **(Supplementary Fig. S8A)**. In the kidney, CisPt caused a pronounced increase in the mRNA levels of *Cdkn1a*, *Cd68*, *Il2ra* and *Il6*, which remained unaffected by *Rac1* deficiency **(Supplementary Fig. S8B)**. Similarly in the liver, CisPt induced the upregulation of *Cdkn1a* and *Mmp3*; while *Cdkn1a* expression was not affected by *Rac1* knockout, *Mmp3* levels were further elevated in *Rac1*-deficient animals **(Supplementary Fig. S8C)**. Taken together, these findings further support the organ-specific roles of RAC1 in the regulation of anticancer drug-induced stress responses, highlighting its importance in modulating preferentially cardiac stress responses to anticancer treatments.

## 4. Discussion

Anticancer drug-induced normal tissue damage impairs the quality of life of cancer patients and limits the effective dose, thereby hampering the therapeutic outcome and prognosis of tumor patients. Therefore, there is a clear clinical need for novel effective and well-tolerated organoprotective measures, which do not interfere with the therapeutic efficacy of the anticancer agents. In this context, pharmacological studies point to RAC1 as a promising target to mitigate anthracycline-induced cardiac injury (71, 72). Although different studies have utilized genetic models to study RAC1 function in specific tissues, comprehensive cross-organ analyses in the context of exposure to different types of anticancer drugs remain limited. We have previously shown that cardiomyocyte-specific *Rac1* knockout using the MHC-Cre system, partially alleviates DOX-induced cardiac injury (57). Moreover hepatocyte-specific *Rac1* deletion by Alb-Cre-mediated knockout, reduces DOX-induced liver DNA damage. However, it remains unclear whether targeting of RAC1 has also the potential (i) to alleviate DOX-induced damage to other organs, especially kidney and liver as major detoxifying organs and (ii) to mitigate organ damage evoked by other classes of anticancer drugs such as the widely used platinating agents. Therefore, in this study, we aimed to figure out whether RAC1 influences anticancer drug-induced normal tissue damage in a cross-organ and cross-agent specific manner. In this way, we wanted to determine whether *RAC1* is a promising therapeutic target for a comprehensive normal tissue protection against various classes of therapeutic agents. To this end we employed a Poly(I:C) inducible *Rac1* knock-out model which enables a broad cross-tissue genetic deletion of *Rac1*.

The results obtained provide evidence that the genetic deletion of *Rac1* confers profound protection against DOX-stimulated acute cardiac, kidney and liver injury as reflected on the levels of DNA damage (i.e., DSB) formation and apoptosis induction. The data demonstrate beneficial cross-organ activity resulting from *Rac1* deficiency in the context of anthracycline treatment. Based on this finding we suggest RAC1 as a highly promising target to prevent multiple adverse effects of anthracycline-based anticancer therapy, particularly in the heart but also in major detoxifying organs. To assess a presumably agent-specific role of *Rac1* in modulating drug-induced DNA damage and apoptosis, we further investigated its effects in response to the platinating agent CisPt. Similar to DOX, CisPt also caused a substantial increase in the percentage of cells harboring DNA damage and undergoing apoptosis in all three organs analyzed. Noteworthy, *Rac1* knockout mitigated CisPt-induced DNA damage in the liver, providing novel evidence of distinct beneficial cross-agent effects of RAC1 regarding the liver. However, *Rac1* knockout neither protected the heart nor the kidney from CisPt-evoked DNA damage induction. Furthermore, *Rac1* knockout did not confer protection against CisPt-triggered apoptosis in any of the organs analyzed. Taken together, these findings highlight the cross-organ beneficial effects of RAC1 knockout in the context of DOX-based anticancer therapy but demonstrate the limitation of RAC1 deficiency for chemoprevention of CisPt-mediated tissue injury.

To gain a more detailed insight into the broad organoprotective effects of *Rac1* knockout regarding DOX-induced DNA damage and apoptosis, we addressed whether DOX treatment leads to cell type-specific responses, particularly in the heart and kidney, which are highly complex organs composed of several different cell types (73, 74). In the heart, DOX caused a significant increase in DNA damage in both cardiomyocyte and non-cardiomyocyte cell fractions. Rac1 knockout reversed this effect in both cell types. Cardiomyocyte apoptosis was not observed after DOX treatment, which is consistent with our previous *in vitro* data (58). However, DOX treatment did stimulate apoptosis in the non-cardiomyocyte fraction, which was significantly reduced in the *Rac1* knockout animals. In the kidney of wild-type animals, DOX treatment induced similarly high levels of DNA damage in cells of both the cortex and the medulla (∼10-fold increase as compared to control), which were almost completely prevented in the absence of RAC1. However, only a moderate increase in the percentage of DNA damage harboring cells (∼3-fold) was found in the glomeruli after DOX treatment, and, furthermore, *Rac1* knockout did not influence the level of DOX-induced DNA damage in glomerular cells. Notably, DOX-induced apoptosis also returned to basal levels in both cortex and medulla cells in the *Rac1* knockout animals. Collectively, these findings demonstrate tissue- and cell-specific responses to DOX, which are differentially modulated by RAC1. These differential RAC1-dependent effects may reflect inherent variations in *Rac1* expression or activity across different tissues, differences in the expression of Top2 isoforms or of factors involved in the regulation of the DDR, DNA repair and/or apoptosis. Moreover, Poly(I:C)-induced knockout efficiency varied markedly across organs. Notably yet, unlike in cisplatin-treated animals, the absence of *Rac1* affected the DOX-induced DNA damage response (DDR) even in organs exhibiting relatively low *Rac1* knockout efficiency, highlighting the distinct role of RAC1 in influencing DOX-induced tissue damage. Summarizing, the results highlight the multifaceted roles of RAC1 in shaping cellular responses to DOX, suggesting that targeting RAC1 holds the therapeutic potential for minimizing especially DOX-induced adverse outcome pathways by conferring organoprotection in a tissue- and cell-type specific manner.

In addition to analyzing the formation of γH2AX foci as surrogate marker of DSB, we investigated the involvement of RAC1 in the regulation of DOX-induced DDR in more detail. To this end, we monitored the protein expression of DDR-associated ATM/ATR-regulated marker proteins such as γH2AX, pKAP1 and pP53. *Rac1* knockout markedly attenuated DOX-induced protein expression of these DDR-related factors both in the kidney and liver, but not in the heart. Under basal situation, pP53 expression was elevated both in the heart and kidney, while basal γH2AX levels were increased only in the heart of *Rac1* knockout mice as compared to wild-type controls. It was previously reported that, in the absence of RAC1, P53 can be activated by MAP kinase-dependent pathways (75). Hence, we suggest that *Rac1* knockout may promote a DSB-independent branch of the DDR in the heart, potentially counteracting its otherwise protective effects against DOX-mediated genotoxic stress. Of note, DOX-induced increase in pP53 levels did not reach statistical significance in the liver, suggesting that DOX treatment activated hepatic DDR-related mechanisms independent of P53. Along these lines, it was previously shown that liver injury strongly activates P21 independently of its upstream regulators P53 and CHK2 (76). Consistent with this, we observed a robust upregulation of *Cdkn1a* mRNA following DOX treatment, which was not mitigated by *Rac1* deletion. Similarly, *Il6* transcript levels were elevated following DOX treatment in the liver independent of the *Rac1* status, suggesting that *Rac1* does not influence these inflammatory and cell cycle regulatory pathways. Interestingly, DOX administration also induced hepatic *Gstm1* mRNA expression, a marker of hepatic detoxification. This response was significantly attenuated in Rac1-deficient animals. This finding suggests that RAC1 deficiency may lower the extent of DOX-induced stress responses of the liver (77).

Addressing cardiac-specific injury markers, the DOX-induced increase in the mRNA levels of *Nppa* was modestly reduced by *Rac1* knockout. We observed a similar trend when calculating the platelet to lymphocyte ratio, which is proposed to be a valuable prognostic marker for cardiac injury (68, 69). These findings suggest that *Rac1* deletion partially alleviates the expression of systemic indicators of DOX-induced cardiac damage. Noteworthy, DOX treatment led to a pronounced increase in the mRNA levels of *Cdkn1a*, *Ctgf*, *Il6* and *Mmp3* in the heart. Importantly, *Rac1* knockout effectively counteracted the DOX-stimulated induction of all of these genes. Interestingly, comparable transcriptional responses were observed in the kidney and liver, as well as following CisPt treatment, pointing to a conserved pattern of stress response activated across different tissues and anticancer agents. However, a marked impact of the *Rac1* status was observed predominantly in the heart, indicating an organ-specific function of RAC1-regulated signaling in fine-tuning cardiac stress responses induced by anticancer agents.

Furthermore, DOX treatment increased the percentage of proliferating (i.e., PCNA positive) cells in the heart, and this effect was further promoted in the absence of RAC1. In contrast, no changes in the percentage of PCNA positive cells were detected in the kidney or liver following DOX treatment. The concurrent emergence of the expression of senescence-associated markers (e.g., *Cdkn1a (p21), Il6, Icam1*) with pro-fibrotic responses (e.g., *Il6, Mmp3*) and increased proliferation in response to DOX exposure, suggests that complex pathophysiological processes are taking place in the heart. Considering the multi-cellular composition of the heart, it is plausible that DOX evoked distinct responses in different cardiac cell populations (58, 74). In this context, it was previously reported that DOX can trigger either apoptosis or senescence in cardiomyocytes, depending on the dose applied (78). Given that cardiomyocyte apoptosis was not observed under our experimental conditions, we speculate that DOX may have triggered senescence in this cell type under our experimental conditions. It is well established that endothelial cells and fibroblasts undergo proliferation in case of injury in order to promote wound healing and repair. However, excessive or dysregulated proliferation may favor the development of fibrosis (79). Yet, investigating αSMA protein levels as an indicator of fibrosis, DOX treatment did not cause any alterations in αSMA expression in any of the organs examined. This is likely due to the early time point of our analysis, as fibrosis is usually a late-onset response to injury (80). Noteworthy in this context, basal αSMA levels in the heart and liver of the *Rac1* knockout animals were significantly higher than in the wild-type controls. These findings indicate that, in the absence of an exogeneous insult, RAC1 signaling may contribute to the maintenance of tissue homeostasis by restraining spontaneous mesenchymal activation. This hypothesis gains support by previous report showing that RAC1 plays an inhibitory role in the TGFβ-induced αSMA expression (81). This phenomenon has been extensively discussed in the context of tumor progression, where RAC1 seems to promote progression and invasiveness via the TGFβ pathway (82). Hence, we speculate that RAC1 may play a dual role in regulating tissue homeostasis, depending on the tissue type, physiological versus pathophysiological conditions, and the presence of exogenous stimuli. These multiple functions make RAC1 a promising target to protect healthy tissues without interfering with the therapeutic efficacy of anticancer drugs.

Collectively, the results of our study provide novel genetic evidence indicating that RAC1-regulated signaling plays a prominent role in the regulation of anthracycline-induced acute injury to both the heart, kidney and liver **(Fig. 7, graphical abstract)**. Rac1 deficiency markedly reduced drug-induced DNA damage formation, induction of apoptosis, as well as the expression of different types of stress markers across all organs following DOX treatment. In contrast, Rac1 deficiency only partially protected against CisPt-induced normal tissue toxicity, with significant effects limited to the heart and liver. This suggests that the protective outcome resulting from *Rac1* deletion is not only organ-but also drug-specific. Taken together, we propose that targeting RAC1 may be a promising approach to alleviate life threatening and life quality impairing adverse effects resulting from anthracycline-based anticancer treatment across different organs, whereas the role of *Rac1* in the context of CisPt-induced normal tissue toxicity is much more limited and warrants further investigation.

**Fig 7.**
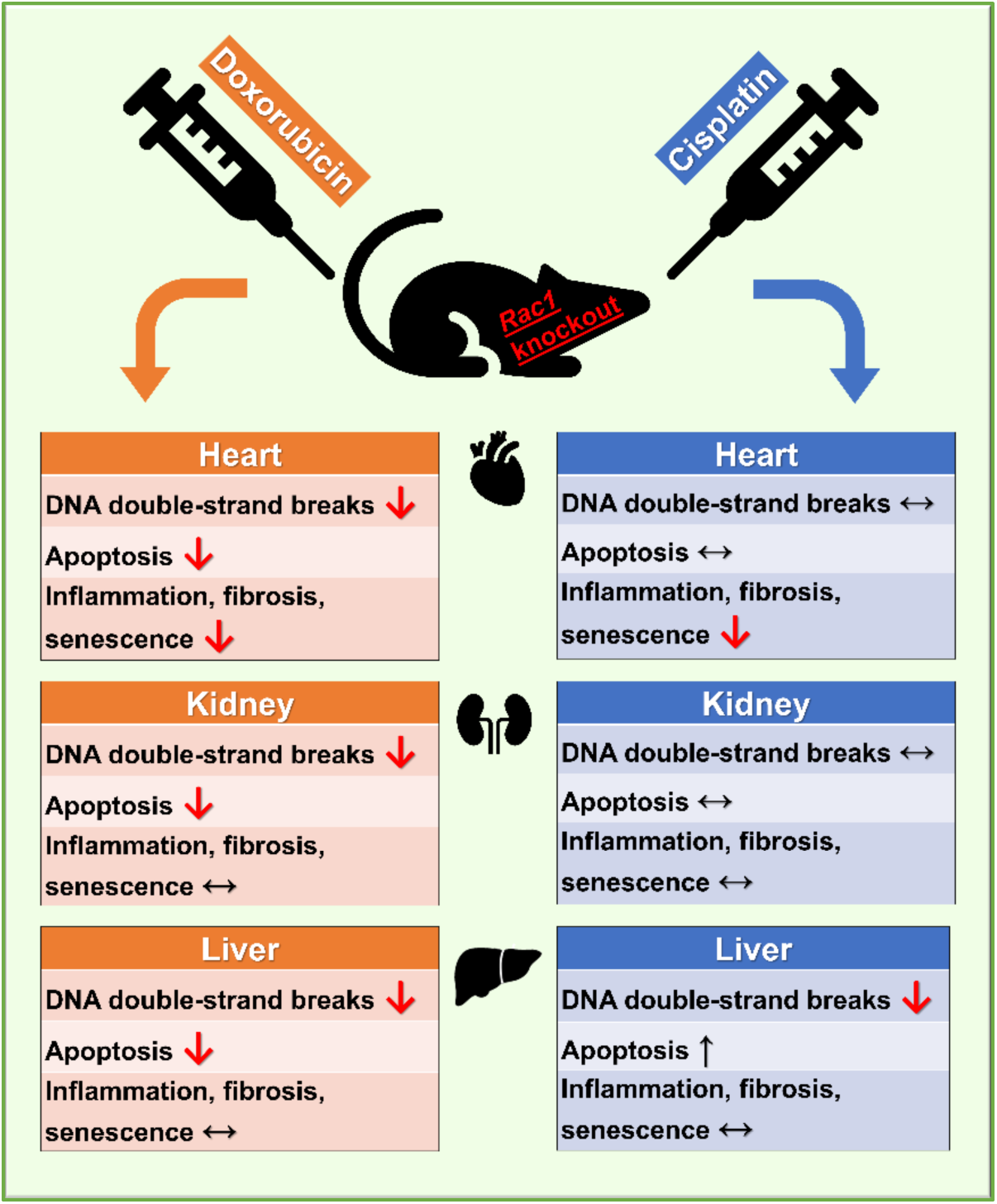
Graphical abstract illustrating the hypothetical role of RAC1 in anticancer therapy-induced normal tissue damage. Our data show that genetic deletion of *Rac1* confers robust geno- and cytoprotection against doxorubicin-induced damage particularly in the heart, as well as in the major detoxifying organs kidney and liver. *Rac1* deletion also shows limited protection against cisplatin-induced injury, with substantial differences regarding organ specificity and adverse outcome pathways affected.

## Statements and Declarations

### Ethics approval

Animal experiments were conducted in accordance with the European Guidelines for the Care and Use of Laboratory Animals (2010/63/EU) and were approved by the North Rhine-Westphalia State Agency for Nature, Environment and Consumer Protection (81-02.04.2020.A140).

### Author contributions

**PK:** Formal analysis, Investigation, Methodology, Validation, Visualization, Writing – original draft, Writing – review & editing.

**SG:** Investigation, Methodology, Validation, Visualization

**JA:** Investigation, Methodology, Validation, Visualization

**CH:** Formal analysis, Supervision, Writing – review & editing

**CH:** Investigation, Visualization

**AV:** Investigation, Visualization

**GF:** Conceptualization, Formal analysis, Funding acquisition, Project administration, Resources, Supervision, Writing – original draft, Writing – review & editing.

### Data availability

No data was used for the research described in the article.

## Acknowledgements

We would like to thank Claudia Gavranic and Lena Abbey (Institute of Toxicology, Düsseldorf, Germany) for excellent technical support.

## Abbreviations

ATM: ataxia telangiectasia mutated
ATR: ATM and RAD3-related
αSMA: alpha smooth muscle actin
Bax: Bcl2-associated X
Bcl2: B-cell lymphoma-2
BW: body weight
Cdkn1a: cyclin dependent kinase inhibitor 1A
CHF: congestive heart failure
CisPt: cisplatin
cTnI: cardiac troponin I
Ctgf: connective tissue growth factor
DAPI: 4’,6-diamidino-2-phenylindole
DDR: DNA damage response
DOX: doxorubicin
DSB: DNA double-strand breaks
EMT: epithelial-mesenchymal transition
FasR/FasL: Fas cell surface death receptor/ Fas ligand
GAPDH: glyceraldehyde-3-phosphate dehydrogenase
GEF: guanine nucleotide exchange factor
GOT: glutamate oxaloacetate transaminase
GPT: glutamate pyruvate transaminase
γH2AX: phosphorylated histone H2AX (serine 139)
Gstm1: glutathione S-transferase Mu 1
iNOS: inducible nitric oxide synthases
Il6: interleukin 6
KAP1: KRAB-associated protein-1
KIM-1: kidney injury molecule-1
KO: knockout
Mmp3: matrix metalloproteinase-3
Nppa/b: natriuretic peptide A/B
PCNA: proliferating cell nuclear antigen
Poly I:C: polyinosinic:polycytidylic acid
RAC1: RAS-related C3 botulinum toxin substrate 1
RHO: RAS-homologous
ROCK: RHO-associated coiled-coil kinase
ROS: reactive oxygen species
TGFβ: transforming growth factor beta
TOP2: topoisomerase II
TUNEL: terminal deoxynucleotidyl transferase dUTP nick end labelling
WGA: wheat germ agglutinin
wt: wild-type

