## Supplementary data for "*Mx1-Cre*-mediated *Rac1* knockout confers multiple protective effects against anthracycline-induced acute normal tissue injury"

**Supplementary Table 1: DNA primers used for genomic PCR analyses.**

| Gene | Forward primer | Reverse primer |
|---|---|---|
| WT *Rac1* | GTGCCAAGGACAGTGACAAG | GGCTCATGAATGCAGAGTCG |
| KO *Rac1* | GTGCCAAGGACAGTGACAAG | GCAATGACAGATGTTCCGCA |

**Supplementary Table 2: Primers used for RT-qPCR mRNA expression analyses.**

| NCBI reference sequence | Gene | Forward primer | Reverse primer |
|---|---|---|---|
| NM_007392.3 | *Acta2* | CATTGCTTCCTCCTCCTC | CTTGGCTTCCTCATCTGAT |
| NM_007393 | *Actb* | GCATTGCTGACAGGATGCAG | CCTGCTTGCTGATCCACATC |
| NM_007527.3 | *Bax* | CTGGACACTGGACTTCCT | GCCACAAAGATGGTCACT |
| NM_009741 | *Bcl2* | GTGTGGTTGCCTTATGTAT | GTATATCCGCTACAAGTTACA |
| NM_009764 | *Brca1* | TTGTGAGCGTTTGAATGA | ACCTGGCTTAGTTACTGT |
| NM_009844.3 | *Cd19* | ATGCCATCTCCTCTCCCTGT | TGCCTCCCTCTTCTACCTCC |
| NM_001291058.1 | *Cd68* | GCCTCATCTCTTCACCAT | AATCTCTTCATTGTGTTCCTAA |
| NM_007658 | *Cdc25a* | TCAAATGAAAGTGAATCAGGAAAT | CTTCATATTCTCGCCATCCA |
| NM_009864.2 | *Cdh1* | CCACAGCCTCATATCATCA | GTATTCGCCAATCTCTAAGTC |
| NM_007669 | *Cdkn1a* | ACCTGAATAGCACTTTGGAAA | TCTGAGCAATGTCAAGAGTC |
| NM_010217 | *Ctgf* | CAAAGCAGCTGCAAATACCA | GGCCAAATGTGTCTTCCAGT |
| NM_011339.2 | *Cxcl15* | CTAGGCATCTTCGTCCGTCC | TTCACCCATGGAGCATCAAG |
| NM_007987.2 | *Fasr* | AGAACCTCCAGTCGTGAA | ATCTATCTTGCCCTCCTTGA |
| NM_010177 | *Fasl* | CTGGAATGGGAAGACACATAT | TGGTCAGCACTGGTAAGA |
| NM_008006.2 | *Fgf2* | GGCTGCTGGCTTCTAAGTGT | CTGTCCAGGTCCCGTTTTGG |
| NM_007836 | *Gadd45a* | GTCGCTACATGGATCAGTG | GTGACTGCTTGAGTAACTACA |
| NM_008084 | *Gapdh* | TCTCCTGCGACTTCAACA | TCTCTTGCTCAGTGTCCTT |
| NM_010358 | *Gstm1* | ACACAGCCTTCATTCTCC | AATTCTAGGAAGCGTGAGTT |
| NM_010442 | *Hmox1* | CCAGAGTCCCTCACAGAT | CCCAAGAGAAGAGAGCCA |
| NM_010493 | *Icam1* | TGCTCAGGTATCCATCCAT | GGAAACGAATACACGGTGAT |
| NM_008361.3 | *Il1b* | CAGCAGCACATCAACAAG | CAGCAGGTTATCATCATCATC |
| NM_008367.3 | *Il2ra* | GCGGATGGGAATCACAAAGC | CGTTAGGTGAATGCTTGGCG |
| NM_031168 | *Il6* | AGTTGCCTTCTTGGGACTGA | CAGAATTGCCATTGCACAAC |
| NM_134249.5 | *Kim-1* | ATGTCACTTCCCCCATCTTG | TGTGTGCCTGGTCCATAGTC |
| NM_001285921.1 | *Mfn2* | ATGTTACCACGGAGCTGGAC | AACTGCTTCTCCGTCTGCAT |
| NM_010809 | *Mmp3* | GCTGTGGGAAAGTCAATGA | GCCATAGTAGTTTTCTAGGTATT |
| NM_010902.5 | *Nfe2l2* | GCAACTCCAGAAGGAACAGG | AGGCATCTTGTTTGGGAATG |
| NM_008725 | *Nppa* | CCTAAGCCCTTGTGGTGTGT | CAGAGTGGGAGAGGCAAGAC |
| NM_008726 | *Nppb* | CTGAAGGTGCTGTCCCAGAT | CCTTGGTCCTTCAAGAGCTG |
| NM_008904 | *Ppargc1a* | CCGAGAATTCATGGAGCAAT | TTTCTGTGGGTTTGGTGTGA |
| NM_009007.2 | *Rac1* | CCTATCATCCTCGTGGGGAC | GGTAGGTGATGGGAGTCAGC |
| NM_011234 | *Rad51* | CAGCGATGTCCTAGATAATGTAG | TTACCACTGCGACACCAA |
| NM_016802 | *Rhoa* | AAGTCTGGGTGCCTCA | AATAATCGTGGTTGGCTTCTAA |
| NM_007483.3 | *Rhob* | CCGAGGTAAAGCACTTCTGC | CCGAGCACTCGAGGTAGTCA |
| NM_001291859.1 | *Rhoc* | TCACAGGAGGAGTCTAGG | CCCACCAAGTTAAGTCAAAT |

| | | | |
|---|---|---|---|
| NM_011577.1 | *Tgfb1* | CAACAACGCCATCTATGAG | AAGGTAACGCCAGGAATT |
| NM_011623 | *Top2a* | CTTCAGGAGCCGTCACCAT | GAGCAGTATATGTTCCAGTTGT |
| NM_011640 | *Trp53* | AAGTTCTGTAGCTTCAGTTCAT | GGCAGTCATCCAGTCTTC |
| NM_011691.4 | *Vav1* | AGATTGATGACACCGCAGAG | CGCATTAGGTCCTCGTAGATC |
| NM_009500.2 | *Vav2* | CATTGAGAAGAACTACATGGG | CCTTAAACTCCAGAAAGACC |
| NM_011701.4 | *Vim* | CGCAAGATAGATTTGGAATAGA | CAGTAACAAGTTGGTCAGATAT |
| NM_001355058.1 | *Wee1* | GTAGTCTCTATTCATGGACACA | GTTGCTTTCAGTAATTGTAATTCTTT |
| NM_028012 | *Xrcc4* | TGCCTGGACACCATTACA | CTTCTCATTCAGCACCAAGAT |

### Supplementary Figures

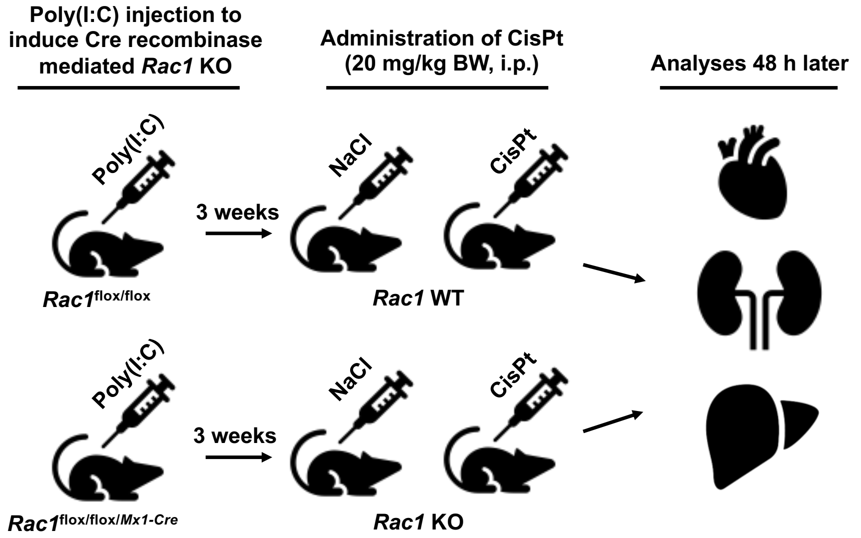


**Supplementary Figure S1: Generation of *Cre*-recombinase-mediated *Rac1* knockout mice and CisPt treatment.**

Poly(I:C) was administered to *Rac1*$^{\text{flox/flox/}Mx1\text{-}Cre}$ and *Rac1*$^{\text{flox/flox}}$ mice to induce a *Cre*-recombinase-mediated *Rac1* knockout (KO) and matched *Rac1* wild-type controls (WT), respectively. A single injection of CisPt (20 mg/kg BW, i.p.) was administered 3 weeks later and analyses were performed after post-incubation period of 48 h. Control groups were treated with saline.

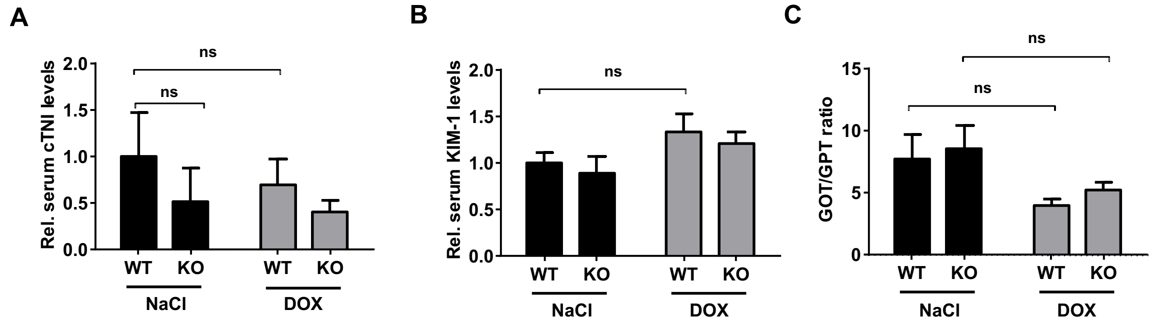


**Supplementary Figure S2: Influence of DOX treatment on serum levels of organ toxicity-related markers in *Rac1* WT and *Rac1* KO mice.**

Serum samples were analysed for the cardiac damage marker cTNI, kidney damage marker KIM-1 and liver damage markers GOT and GPT. **(A)** Relative cTNI levels in saline-treated *Rac1* WT control group are set to 1.0. **(B)** Relative KIM-1 levels in saline-treated *Rac1* WT control group are set to 1.0. **(C)** GOT/GPT ratio was calculated. Data represent the mean + SEM from n=3-8 mice per experimental group. ns, not significant, two-tailed Student's *t*-test.

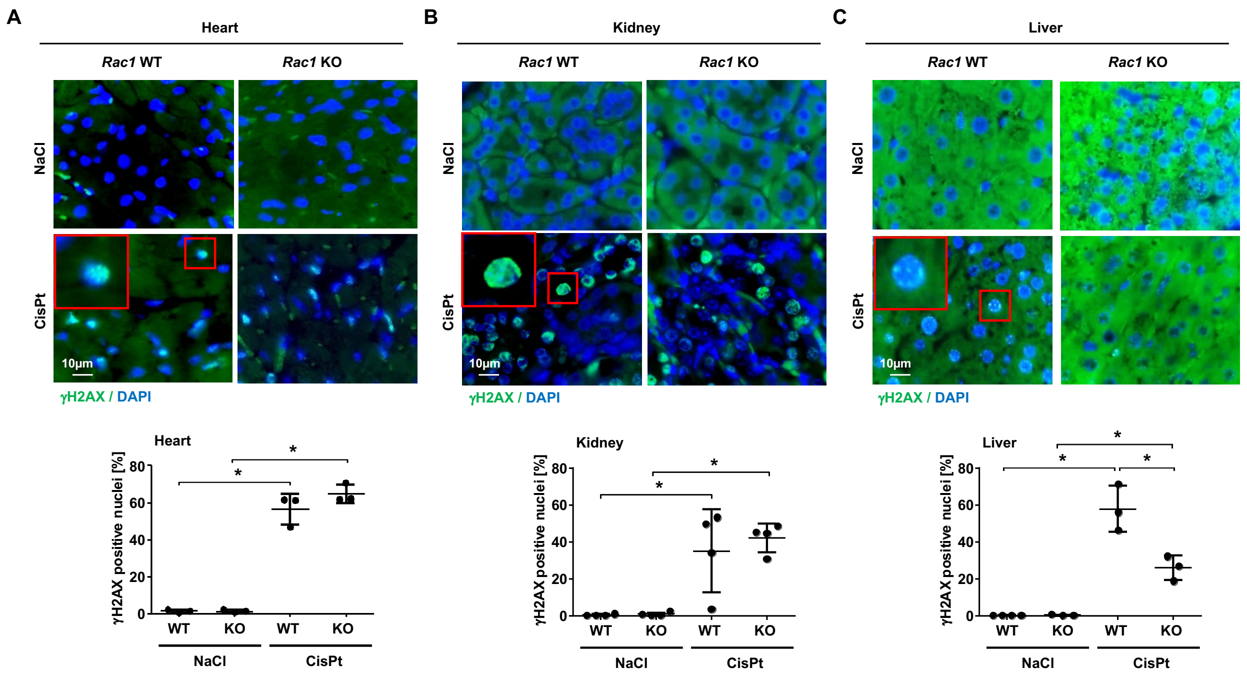


**Supplementary Figure S3: Influence of *Rac1* knockout on CisPt-induced DSB levels in the heart, kidney and liver.**

*Rac1* WT and *Rac1* KO mice were treated with saline or CisPt (20 mg/kg BW) as shown in Supplementary Fig. S1.

**A-C)** DNA DSB were visualized by immunohistochemical staining of γH2AX in the heart **(A)**, kidney **(B)** and liver **(C)** tissue sections. The percentage of γH2AX positive nuclei was quantified. Data shown are the mean ± SD from n=3-4 animals per experimental group with two technical replicates and >150 nuclei being analyzed per condition. *$p \leq 0.05$; two-tailed Student's *t*-test. The upper panels show representative pictures. γH2AX (green), DAPI (blue); 100x objective.

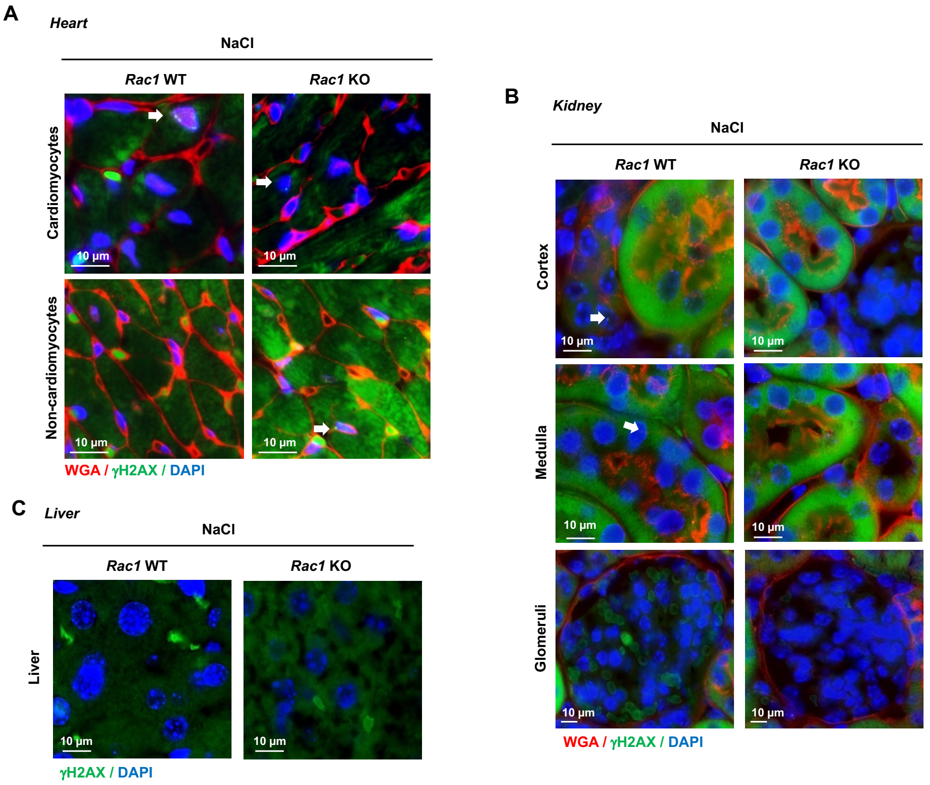


**Supplementary Figure S4: Representative pictures obtained from saline-treated control groups.** **A-C)** DNA DSB were visualized by immunohistochemical staining of γH2AX in the heart **(A)**, kidney **(B)** and liver **(C)** tissue sections. WGA was used to visualize cell borders in the heart **(A)** to distinguish between cardiomyocytes and non-cardiomyocytes and in the kidney **(B)** to distinguish between cortex, medulla and glomeruli. WGA (red), γH2AX (green), DAPI (blue); 100x objective. White arrows indicate γH2AX positive nuclei.

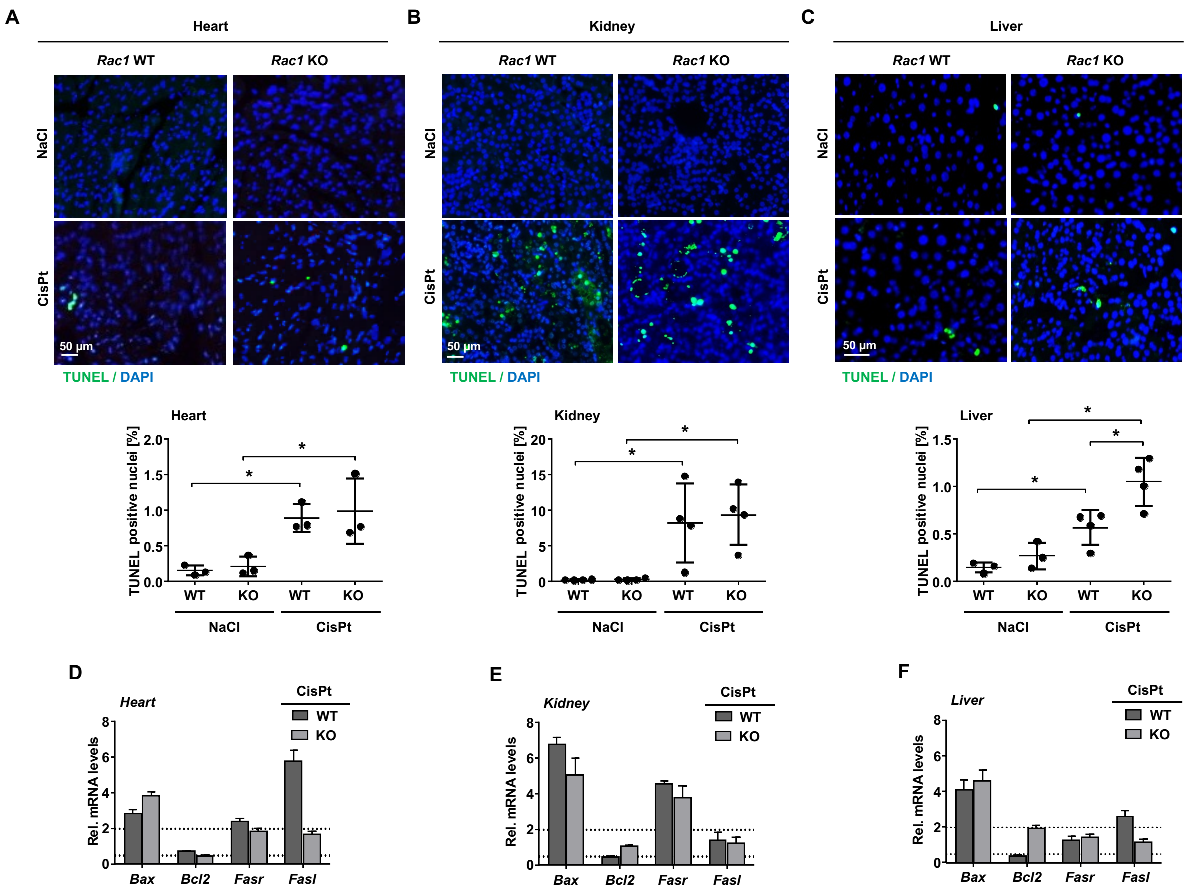


**Supplementary Figure S5: Influence of *Rac1* knockout on CisPt-induced apoptosis in the heart, kidney and liver.**

*Rac1* WT and *Rac1* KO mice were treated with saline or CisPt (20 mg/kg BW) as shown in Supplementary Fig. S1.

**A-C)** Apoptotic cells were visualized by immunohistochemical TUNEL staining in the heart **(A)**, kidney **(B)** and liver **(C)** tissue sections. The percentage of TUNEL positive cells was quantified. Data shown are the mean ± SD from n=3 animals per experimental group with two technical replicates and >300 cells being analyzed per condition. *$p \leq 0.05$; two-tailed Student's *t*-test. The upper panels show representative pictures. TUNEL (green), DAPI (blue); 20x objective.

**D-F)** mRNA expression levels of genes involved in the regulation of apoptosis were detected by RT-qPCR in the heart **(D)**, kidney **(E)** and liver **(F)** tissues. mRNA expression levels were normalized to *Gapdh* and *Actb* as described in methods. Relative mRNA expression in saline-treated *Rac1* WT control group was set to 1.0 and relative mRNA levels were shown as mean + SEM from triplicate determinations using pooled RNA samples isolated from n=3 animals per experimental group.

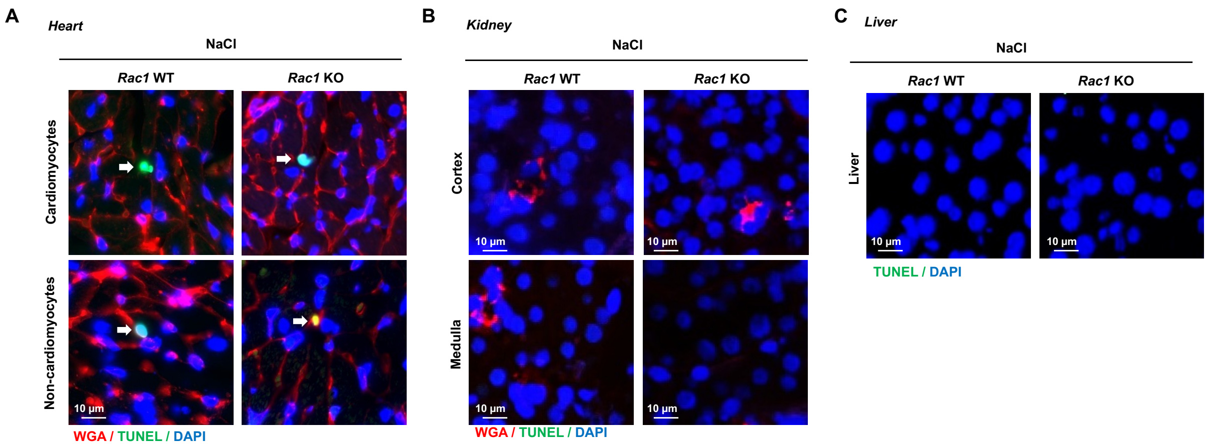


**Supplementary Figure S6: Representative pictures obtained from saline-treated control groups. A-C)** Apoptotic cells were visualized by immunohistochemical TUNEL staining in the heart **(A)**, kidney **(B)** and liver **(C)** tissue sections. WGA was used to visualize the cell borders in the heart **(A)** to distinguish between cardiomyocytes and non-cardiomyocytes and in the kidney **(B)** to distinguish between cortex and medulla. WGA (red), TUNEL (green), DAPI (blue); 100x objective. White arrows indicate TUNEL positive nuclei.

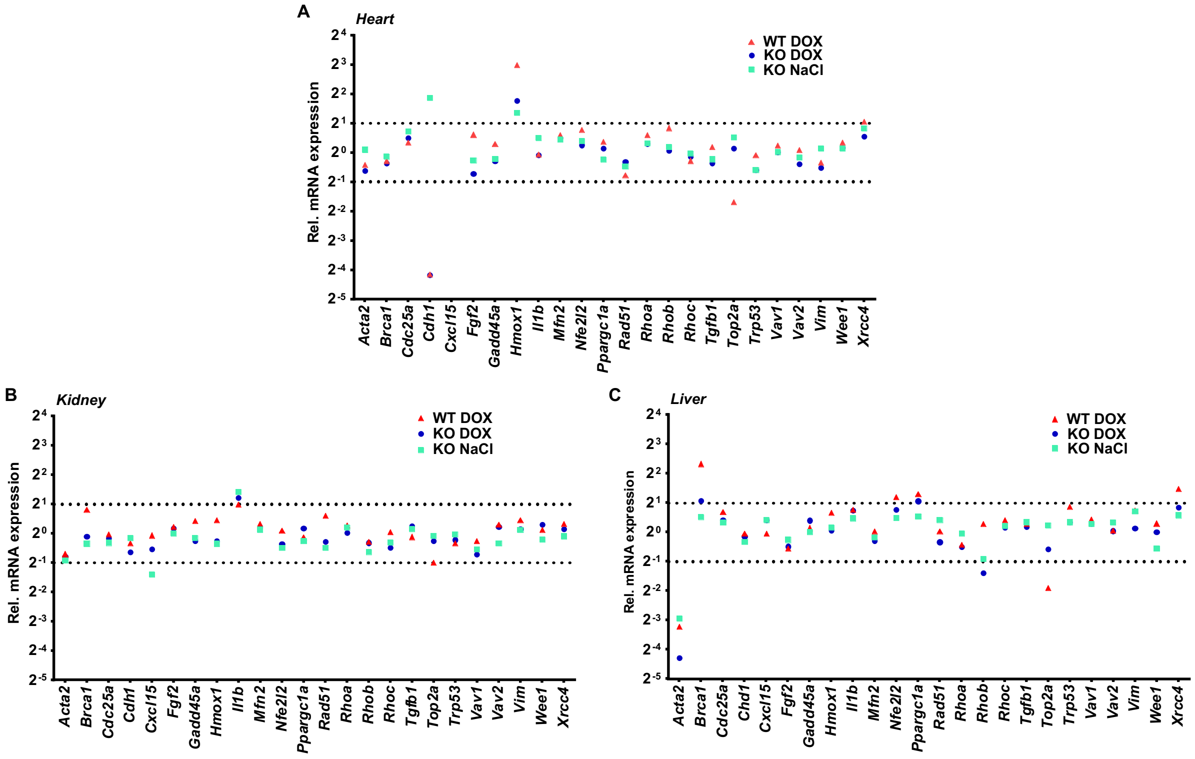


**Supplementary Figure S7: Influence of *Rac1* knockout on basal and DOX-induced alterations in the mRNA expression of genotoxin susceptibility-related genes in the heart, kidney and liver.** *Rac1* WT and *Rac1* KO mice were treated with saline or DOX as shown in Fig 1A.

**A-C)** mRNA levels of selected genes involved in the regulation of Ras-homologous (RHO) GTPases, DNA repair, cell death, oxidative metabolism, mitochondrial homeostasis, topoisomerases, senescence, inflammation and fibrosis were detected by RT-qPCR in the heart **(A)**, kidney **(B)** and liver **(C)** tissues. mRNA expression levels were normalized to *Gapdh* and *Actb.* Relative mRNA expression in saline-treated *Rac1* WT control group was set to 1.0 and relative mRNA levels from triplicate determinations using pooled RNA samples isolated from n=3 animals per experimental group are shown.

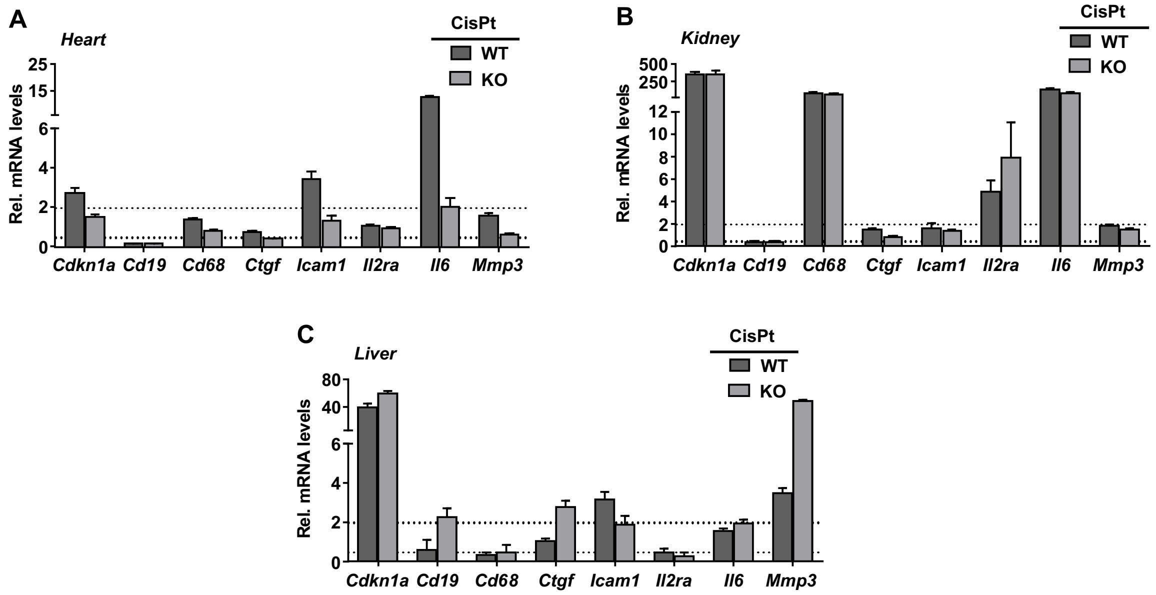


**Supplementary Figure S8: Influence of *Rac1* knockout on CisPt-induced alterations in the mRNA expression of inflammation, fibrosis and senescence-related genes in the heart, kidney and liver.** *Rac1* WT and *Rac1* KO mice were treated with saline or CisPt (20 mg/kg BW) as shown in Supplementary Fig. S1.

**A-C)** mRNA levels of selected genes involved in the regulation of senescence (*Cdkn1a*), inflammation (*Cd19*, *Cd68*, *Icam1*, *Il2ra*, *Il6*) or fibrosis (*Ctgf, Mmp3*) were investigated by RT-qPCR in the heart **(A)**, kidney **(B)** and liver **(C)** tissues. mRNA expression levels were normalized to *Gapdh* and *Actb.* Relative mRNA expression in saline-treated *Rac1* WT control group was set to 1.0 and relative mRNA levels were shown as mean + SEM from triplicate determinations using pooled RNA samples isolated from n=3 animals per experimental group.
